# Adaptive Plasticity Tumor Cells Modulate MAPK-Targeting Therapy Response in Colorectal Cancer

**DOI:** 10.1101/2025.01.22.634215

**Authors:** Oscar E. Villarreal, Yixin Xu, Ha Tran, Annette Machado, Dionne Prescod, Amanda Anderson, Rosalba Minelli, Mike Peoples, Alejandro Hernandez Martinez, Hey Min Lee, Chi Wut Wong, Natalie Fowlkes, Preeti Kanikarla, Alexey Sorokin, Jumanah Alshenaifi, Olu Coker, Kangyu Lin, Chris Bristow, Andrea Viale, John Paul Shen, Christine Parseghian, Joseph R. Marszalek, Ryan Corcoran, Scott Kopetz

## Abstract

MAPK pathway inhibitors (MAPKi) are increasingly used in the treatment of advanced colorectal cancer, but often produce short-lived responses in patients. Although acquired resistance by *de novo* mutations in tumors have been found to reduce response in some patients, additional mechanisms underlying the limited response durability of MAPK targeting therapy remain unknown. Here, we denote new contributory tumor biology and provide insight on the impact of tumor plasticity on therapy response. Analysis of MAPKi treated patients revealed activation of stemness programs and increased ASCL2 expression, which are associated with poor outcomes. Greater ASCL2 with MAPKi treatment was also seen in patient-derived CRC models, independent of driver mutations. We find ASCL2 denotes a distinct cell population, arising from phenotypic plasticity, with a proliferative, stem-like phenotype, and decreased sensitivity to MAPKi therapy, which were named adaptive plasticity tumor (APT) cells. MAPK pathway suppression induces the APT phenotype in cells, resulting in APT cell enrichment in tumors and limiting therapy response in preclinical and clinical data. APT cell depletion improved MAPKi treatment efficacy and extended MAPKi response durability in mice. These findings uncover a cellular program that mitigates the impact of MAPKi therapies and highlights the importance of addressing tumor plasticity to improve clinical outcomes.

## Introduction

Treatment of colorectal cancer (CRC), particularly in the context of metastatic disease, continues to be a major clinical challenge due to heterogeneity and therapy resistance^1–4^. As the genetic landscape of CRC has been uncovered, use of targeted therapies has increased for metastatic CRC (mCRC), primarily in the use of MAPK pathway inhibitors (MAPKi)^5–9^. Several MAPKi containing regimens have now been approved including the use of first line anti-EGFR (EGFRi) agents combined with chemotherapy, and second line use of EGFRi with BRAF inhibitors (BRAFi) for patients with BRAF^V600E^ mutated tumors, or EGFRi with KRAS G12C inhibitors (G12Ci) for use in those with KRAS^G12C^ mutations^10–13^. However, results from clinical trials using MAPKi therapies repeatedly demonstrate patients obtain short-lived treatment responses, as noted by typical progression free survival (PFS) of 4-6 months^14–16^. MAPKi therapy in patients with other cancers, such as melanoma and NSCLC, produce longer responses with PFS ranging 10-15 months, suggesting the presence of unique tumor biology limiting the durability of MAPKi therapy in CRC, which suppresses outcome improvement for these patients^17–20^. While tumor cell acquisition of *de novo* mutations has been shown to be a factor contributing to shorter MAPKi therapy response in patients, additional mechanisms remain elusive and warrant investigation given the need for therapy optimization and the potential clinical impact^21–24^.

Many studies have reported on the important effects of tumor stem-like cells and plasticity on the initiation and progression of CRC, as well as its response to typical treatments^25–29^. While multiple CRC stem cell phenotypes have been reported, studies have shown chemotherapy enriches stem-like cells, and recent works have uncovered the central role of plasticity on metastasis and how its impacted by chemotherapy^30–35^. However, little is known regarding the impact of MAPKi therapy on tumor cell stemness or plasticity, and the effect of these specialized cell populations on MAPKi therapy response in patients with CRC.

To identify treatment related molecular pathway alterations that impact response, we used sequencing data and clinical outcomes from MAPKi-treated patients with CRC, which showed treatment activated stem-like programs in tumor cells. Combining transcriptomic datasets, clinical trial data, patient-derived models, cellular barcoding, and clinically relevant MAPKi therapies, we sought to delineate the underlying tumor biology limiting response durability in CRC. Expression of the transcription factor ASCL2, which denotes a cell population with stem activity in CRC^36^, increased with MAPKi treatment in both patient and preclinical samples. Transcriptomic and functional profiling revealed ASCL2^+^ tumor cells as proliferating, stem-like, high plasticity and MAPKi resistant cells that are part of a dynamic state, representing a distinct phenotype we term Adaptive Plasticity Tumor (APT) cells. Inhibition of MAPK signaling produced APT cell phenotype induction, resulting in APT cell enrichment, independent of tumor mutation status or therapy regimen. Finally, concurrent MAPKi therapy and toxin-mediated APT cell depletion improved efficacy and prolonged response durability in CRC models. Together these data show MAPKi therapy limits response durability in mCRC through the induction of a treatment resistant and high plasticity phenotype, termed APT cells, which aligns with other widely recognized progenitor or stem-like cell phenotypes, and importantly demonstrate that elimination of this tumor cell compartment improves MAPKi therapy results.

## Results

### MAPKi therapy activates stem programs in CRC

Investigating the limited response of MAPKi treatment in CRC using patient data is limited due to a lack of data availability. In a recent trial (NCT03668431)^16^, patients (n = 37) with BRAF^V600E^ tumors received combination MAPKi and immunotherapy, and paired scRNAseq data was obtained pre- and post-treatment (**Fig. 1A**). Patients were classified based on PFS as responders (R) and non-responders (NR), enabling use of data to explore therapy-induced transcriptomic changes that impact treatment response. After QC and pseudobulking of epithelial cells, principal component analysis showed clustering of NR (n = 6) and R (n = 5) patient samples (**Fig. 1B**). Genes differentially expressed with treatment (On vs. Pre) between the NR and R group were identified to explore transcriptomic differences impacting response. Stem-related genes, such as SOX9, CDX2, and ALDH1A1, were upregulated while MAPK pathway and proliferation associated genes were downregulated in both patient populations. REG4, C2CD4A, STYL2 and other immune response genes were uniquely enriched in R group, as observed in the original study (**Fig. 1C**). A list of gene signatures (n = 52) of clinical or molecular features reported to affect treatment response or outcomes in CRC was compiled and GSEA conducted to identify modulated pathways that impact therapy response (**Supplementary Table 3**). As expected, among the significantly altered pathways, Immune Response Activation and three other immune-based gene sets were seen, indicating the importance of immune pathways in determining therapy sensitivity (**Fig. 1D**). Interestingly, six independent stemness or differentiation related signatures – including the EpiHR, Revival ISC, and SCS signatures – were significantly modulated, suggesting cancer stem cell biology impacts MAPKi response in patients with CRC.

**Figure 1.**
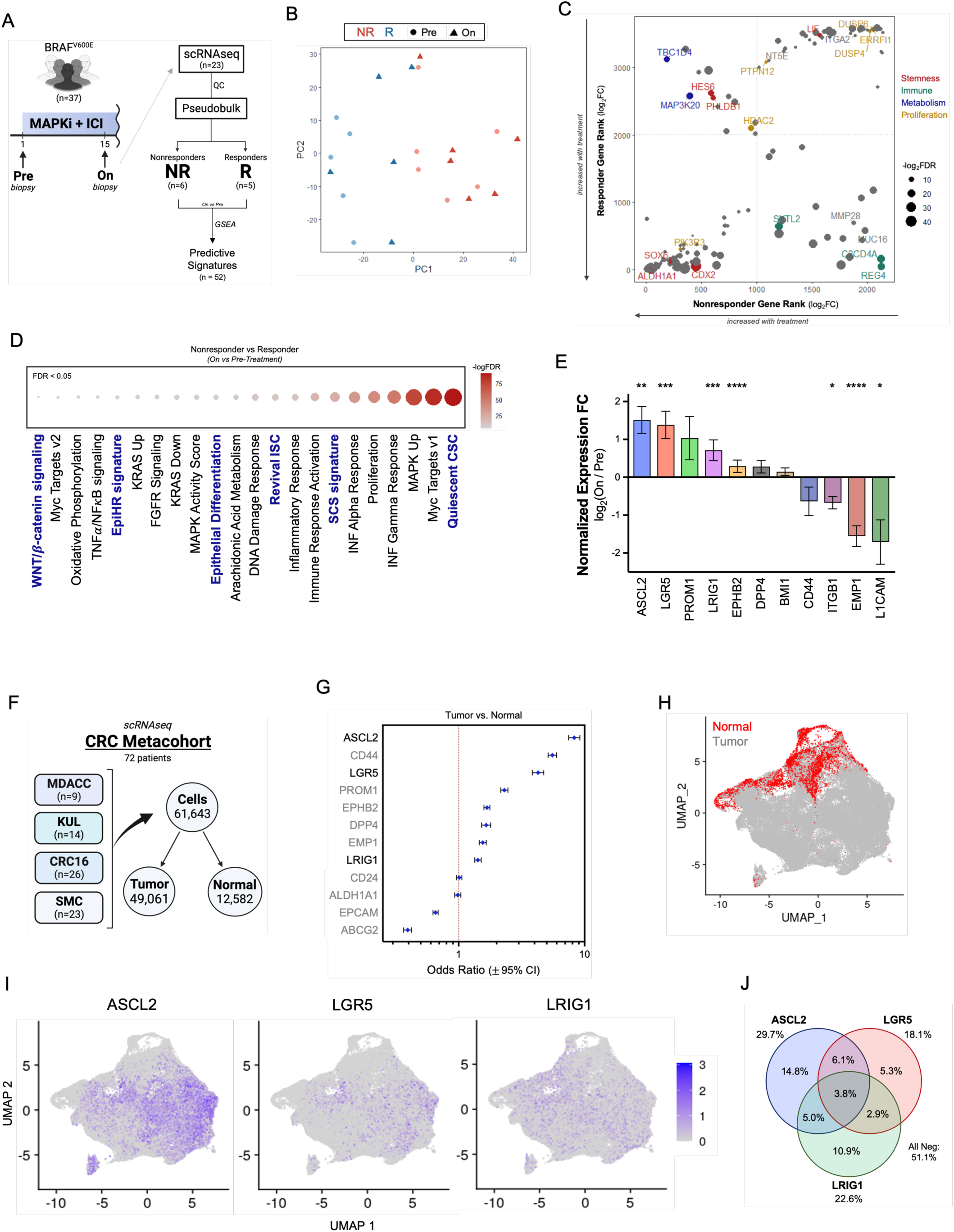
MAPKi treatment increases ASCL2 and stem programs in patients with mCRC. **A)** Outline of BRAFi patient cohort clinical trial (NCT03668431) who received dabrafenib, trametinib and sparatizumab (ICI); paired samples were collected from patients and scRNAseq data processed; pseudobulk analysis used data from 6 nonresponders (NR) and 5 responders (R) patients. **B)** PCA plot of pseudobulked NR (red) and R (blue) samples at pre-treatment (circles) and on-treatment (triangle) timepoints; clustering by response indicates potential shared underlying biology. **C)** Genes significantly altered by therapy, in both NR and R, were ranked by log2FC and DEGs with p<0.05 that appear in both ranked lists are shown. Dotted lines indicate where the ranked genes switch from upregulated (log2FC > 0) to downregulated. Genes relevant to stemness (red), immune processes (green), metabolism or stress response (blue), and proliferation (gold) are labeled by color and size of points depicts corresponding FDR values. **D)** Results from GSEA using DEGs from comparison of treatment changes between NR and R, showing signatures with FDR < 0.05; signature names in blue are stem-related programs. **E)** Change in normalized expression of CRC stem cell markers (On vs. Pre-treatment) per patient (n = 11) and DEG analysis p-adjusted values displayed; *p.adj < 0.05, **p.adj < 0.005, ***p.adj < 0.0005, ****p.adj < 0.0001 (mean ± SD). **F)** CRC Metacohort of scRNAseq data from 72 patients. **G)** Cell expression of stem cell markers in tumor and normal tissue samples; odds ratio and 95% CI for each gene. **H)** Metacohort UMAP colored by tissue sample type, showing tumor (grey) and normal (red) tissue cells. **I)** UMAP showing gene expression of top three stem markers (ASCL2, LGR5, LRIG1). **J)** Venn diagram of cells expressing ASCL2, LGR5, and LRIG1 phenotypes in the tumor cell population; precents calculated based on all tumor cells in dataset.

Multiple markers (CD44, CD133, LRIG1, LGR5) have been reported to denote stem-like cells in CRC but the biological or clinical difference of the cell populations are largely unknown^33,37^. To further determine if MAPKi therapy affects the tumor stem cell compartment, changes in expression of CRC stem cell marker genes induced by treatment were analyzed across all patients. Expression of several markers was impacted by treatment, with ASCL2, LGR5, and LRIG1 showing the greatest significant enrichment in patients treated with MAPKi therapy (**Fig. 1E**). However, expression of stem cell markers used in CRC is often non-overlapping and is not limited to tumor cell populations. In an effort to select the best marker denoting the tumor-specific cell population of interest, a second scRNAseq dataset was compiled using published data^34,38^. The CRC Metacohort contained paired tumor and normal-adjacent tissue samples from 72 patients with CRC, and consisted of 61,643 epithelial tumor or normal cells (**Fig. 1F; Supplementary Fig. 2A-B**). Cells were classified based on CRC stem marker expression and cell number compared between the tumor and normal cell population, which showed ASCL2 had the greatest association with tumor cells (OR = 8.27; 95% CI: 7.45–9.18) compared to other markers (**Fig. 1G**). Between ASCL2, LGR5, and LRIG1, the CRC Metacohort had higher expression of ASCL2, with the greatest expression levels seen in tumor cells (**Fig. 1H, I**). Finally, in the tumor cell population (**Supplementary Fig. 2C**), ASCL2 was more abundantly expressed (29.7% cells) and had greater co-expression with the other markers compared to LGR5 or LRIG1 (**Fig. 1J**). Overall, these results indicate ASCL2 as the best marker for further investigating of the underlying treatment-related and tumor specific stem cell biology observed with MAPKi treatment in CRC.

### ASCL2 in CRC and effect of MAPKi treatment

Commonly reported to be involved in neurogenesis and placental development, the bHLH transcription factor ASCL2 also plays an important role in maintenance and regeneration of intestinal stem cells (ISCs)^39–41^. Some of the molecular mechanisms of ASCL2’s function involves the Wnt, Notch and Hippo (by YAP1 and KLF5) pathways, but its complex signaling dynamics have not been fully uncovered^42–44^. Compared to other primary malignancies CRC tumors displays the highest ASCL2 expression levels in TCGA data (**Supplementary Fig. 1A**). Further, analysis of Human Protein Atlas data showed high ASCL2 mRNA expression is predictive of worse outcomes for patients with mCRC (**Supplementary Fig. 1B**). Together with our findings that MAPKi therapy activates stem programs and increases ASCL2 expression, these data support the clinical relevance of ASCL2 and suggest treatment may activate aggressive CRC biology with poor outcomes.

To further study the observed tumor biology from the clinical data, we tested if MAPKi therapy has the same impact in our preclinical models. Patient-derived xenografts (PDX), organoids (PDO), and cell lines (PDC) from various driver mutation backgrounds were utilized to see if MAPKi therapy alters ASCL2 expression in CRC models (**Supplementary Fig. 1C**). To simulate clinical relevance, MAPKi therapy was tailored to each model based on its driver mutations rather than using a single MAPKi combination (**Supplementary Fig. 1D**). As such, C1047 and B8219 PDO models were treated with MAPKi therapy (**Fig. 2A-B**) along with PDC models (B8219, C1138, C1035) to assess for changes in ASCL2 mRNA expression. Interestingly, all models showed an increase in ASCL2 expression with MAPKi relative to the control, independent of model type or treatment used (**Fig. 2C**). To determine if MAPKi treatment results in protein level changes, ASCL2 IHC staining was performed on MAPKi-treated PDX models. Both B8026 and B1003 displayed increased ASCL2 nuclear staining in treated samples compared to control, indicating greater ASCL2 activity in MAPKi treated tumors (**Fig. 2D**). Additionally, RNAseq data was compiled from previously conducted PDX studies that tested MAPKi therapy combinations with various CRC models. Across all models, MAPK signaling was successfully suppressed by the corresponding treatment, as shown by GSEA comparing MAPKi treated samples versus vehicle using MAPK Pathway Activity Score (MPAS). Comparison of normalized mRNA values between samples revealed higher ASCL2 expression with MAPKi-treatment relative to control in all models, independent of driver mutation or therapeutic regimen (**Fig. 2E**). Therefore, preclinical CRC models recapitulate clinical findings that MAPKi therapy increases ASCL2 and demonstrates this effect is seen widely across multiple CRC models and treatments.

**Figure 2.**
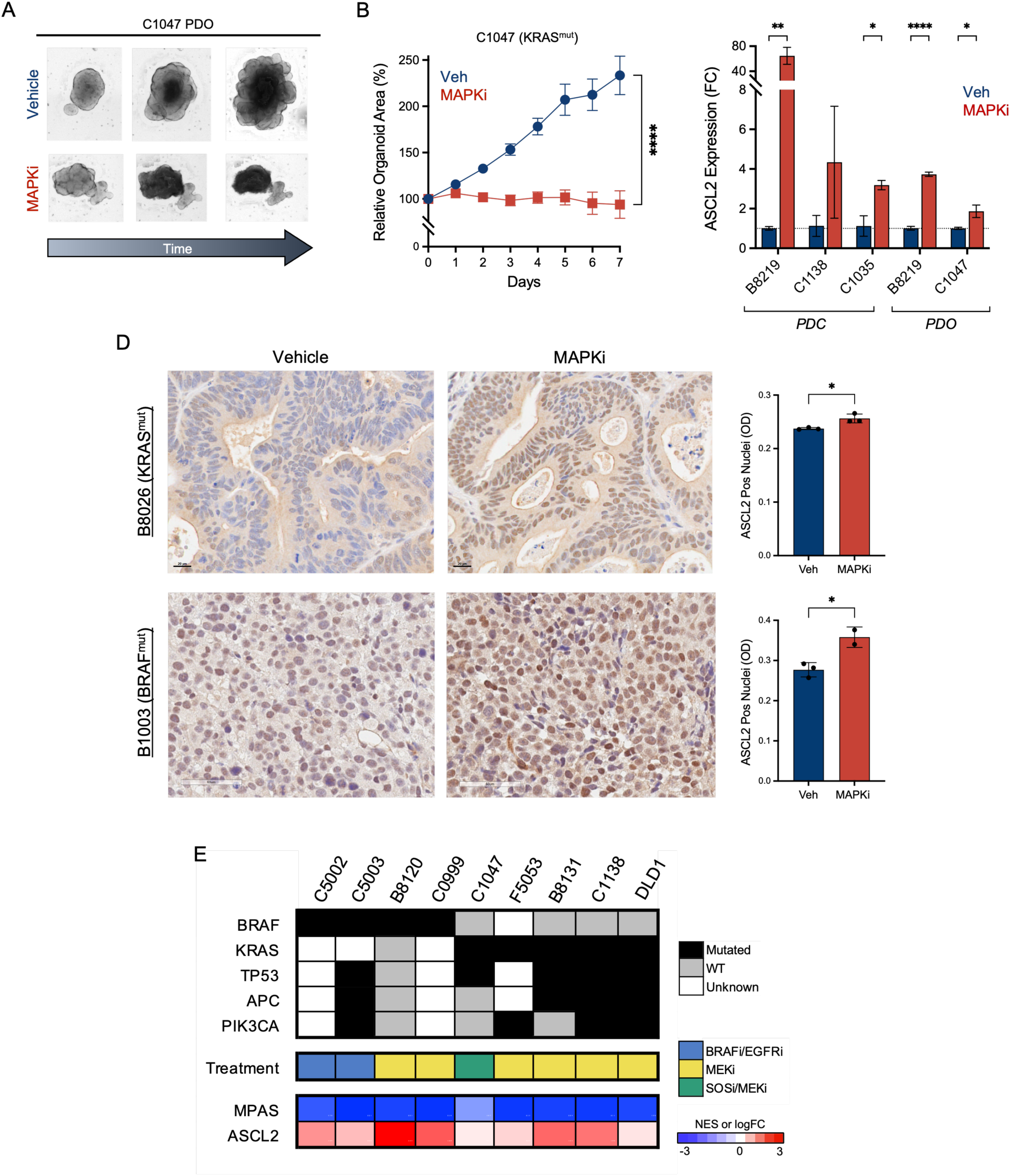
MAPKi increases ASCL2 in preclinical CRC models. **A)** Representative images of KRAS^mut^ C1047 organoids during treatment course with vehicle or MAPKi therapy (BI-3406, trametinib). **B)** C1047 PDO response to vehicle (blue) or MAPKi (red), reported as sum organoid area per well (n = 3) relative to starting point (two-tailed t-test; mean ± SEM). **C)** ASCL2 mRNA expression fold change in PDOs and PDCs after MAPKi (red) treatment relative to vehicle (blue) for B8219 (p = 0.005), C1138, C1035 (p = 0.033), B8219 (p < 0.0001), and C1047 (p = 0.027) models (one-tailed t-test; mean ± SEM). **D)** ASCL2 IHC staining of vehicle or MAPKi treated PDX tumors, and average nuclear optic density (OD) of positively stained cells in B1003 (p = 0.023) and B8026 (p = 0.017) tumors (n = 3) for each treatment group (two-tailed t-test; mean ± SD). **E)** Summary table of MAPKi treated CRC tumors with mutation status and treatment used for each model; RNAseq results for MAPK pathway activity score (MPAS) and ASCL2 expression as log_2_FC of MAPKi treated samples normalized to vehicle (n = 3) for corresponding models.

### ASCL2^+^ phenotype is a stem-like, MAPKi resistant state

Patients with mCRC and patient-derived models that received MAPKi therapy demonstrated elevated ASCL2 expression, and PDX data indicated higher ASCL2 activity with treatment. In intestinal crypts, ASCL2 functions in a Wnt/TCF-dependent manner and serves as the master regulator of LGR5^+^ ISCs, which play important roles in the tumor initiation and metastatic progression of CRC^42,45–47^. Using their STAR (stem cell ASCL2 reporter) minigene system, Oost et al. showed ASCL2 denotes a CRC cell population resembling LGR5^+^ ISCs and containing stem-cell features^36^. While several studies have similarly reported ASCL2^+^ cells display stem cell properties^48,49^, a detailed characterization of CRC ASCL2^+^ cells is needed and would provide valuable insight on the observed changes with MAPKi therapy in CRC.

For improved cell identification, an ASCL2 gene signature was generated to more thoroughly capture cells with the ASCL2^+^ phenotype. Using published GSE99133 data, differential gene analysis of flow-sorted ASCL2^+^ (STAR^+^) and ASCL2^-^ (STAR^-^) cell populations was conducted. Of the resulting DEGs, genes found to be upregulated in ASCL2^+^ cells were used for the ASCL2 gene signature (gsASCL2). To account for the ASCL2^+^ phenotype arising from suppressed gene networks, we also computed an ASCL2 Index, derived from both the gsASCL2 and downregulated gene signature values. ASCL2 signatures were validated with the CRC scRNAseq metacohort, where ASCL2 gene expression significantly correlated with gsASCL2 expression scores (Pearson correlation, p = 0.008), and gsASCL2 successfully denoted a cell population of ASCL2^+^ cells (**Supplementary Fig. 2D-F**). Given the limitations of scRNAseq from dropout events, identification of cell populations is more robust when using a gene signature, therefore we use both ASCL2 and gsASCL2 to classify cells for the subsequent analysis.

As the most comprehensive dataset with samples from 72 patients, the CRC metacohort was used for transcriptional characterization of ASCL2^+^ cells. Tumor cells from each patient were classified as ‘positive’ or ‘negative’ based on ASCL2 and gsASCL2 expression to enable analysis. Since ASCL2^+^ cells are reported to display stem-like features, we used previously published stemness gene signatures to assess the two cell population. Comparison of cells scored with the pan-cancer dedifferentiation Stem Index (miRNAsi)^50^ revealed ASCL2^+^ and gsASCL2^+^ groups have higher mean stemness expression than their respective negative cell populations (**Fig. 3A**). Stem potency, a measure of the extent of stemness in a cell, was computed for each tumor cell using cytoTRACE2, and results showed ASCL2^+^ cell populations have higher stem potency (**Fig. 3B**), confirming that ASCL2^+^ cells have a stem-like phenotype. Of note, in each case the gsASCL2 classified cells showed greater differences than the ASCL2 cells, suggesting the gene signature better captures the stem-like phenotype of interest than ASCL2 expression alone. We next tested if ASCL2^+^ cells represent a population with high plasticity. Distinct from stemness, cellular plasticity is the ability with which a cell can change its phenotype. Although not directly quantifiable, plasticity can be assessed using entropy, which quantifies the number of distinct transcriptomic programs active within a cell. As such, entropy was calculated for all metacohort tumor cells and mean values compared between groups. Both ASCL2^+^ and gsASCL2^+^ cell populations had higher entropy, representing cells in a higher plasticity state than the negative cell groups (**Fig. 3C**). Interestingly, while no association was identified between stem potency and gsASCL2 (**Supplementary Fig. 2D**), a strong correlation (r = 0.8, p = 2.0x10^-15^) was observed when gsASCL2 expression was compared with entropy values (**Fig. 3D**). Together, these results suggest that while ASCL2^+^ cells have a stem-like phenotype, it is cellular plasticity that increases with ASCL2 activity in CRC cells.

**Figure 3.**
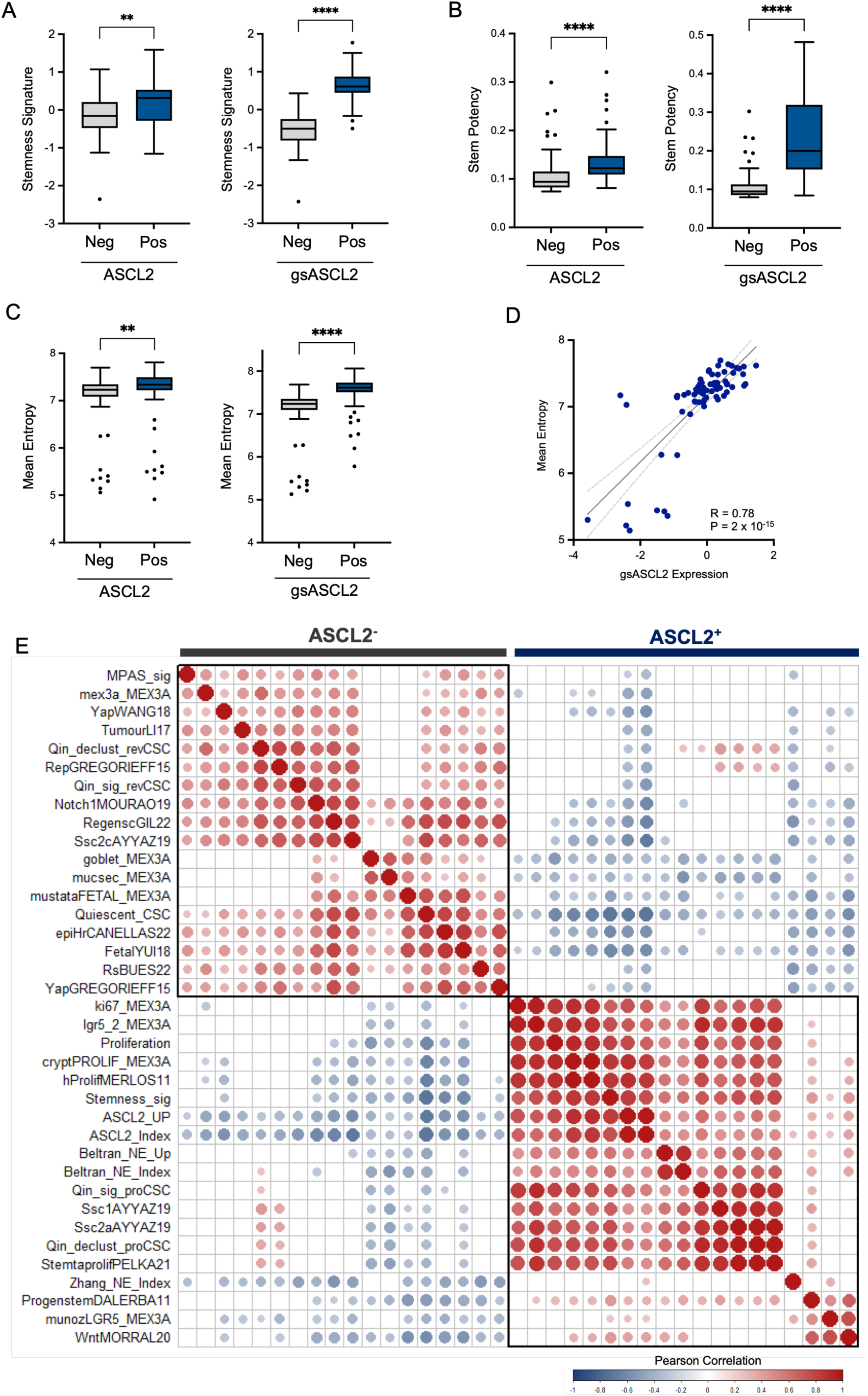
Transcriptomic profiling of ASCL2^+^ cells in CRC Metacohort. **A-C)** CRC metacohort average stemness signature (**A**), stem potency (**B**), and entropy (**C**) per patient (n = 72) from tumor cells classified as negative (grey) or positive (blue) by ASCL2 gene (left) and gsASCL2 signature (right) expression (two-sided Mann-Whitney). Box plots depict median, first and third quartiles, whiskers reflect 1.5*IQR, and outliers shown as points; p ≤ 0.006 (**), p < 0.0001 (****). **D)** Mean tumor cell expression of gsASCL2 and entropy per patient (n = 72), with simple linear regression trendline (black) and 95% CI (dotted grey) depicted (two-sided Pearson correlation). **E)** Pearson correlation plot of stem-cell signatures, including gsASCL2 and ASCL2 Index, using pseudobulked patient data (n = 63); only signatures with significant (p < 0.05) correlations are shows (n = 31), circle size and color indicate correlation coefficient as shown by scale.

With the multitude of stem-like phenotypes identified in CRC tumors, we sought to determine if the ASCL2^+^ cell population expression profile overlaps with any other published phenotype. As such, using more than 75 signatures (**Supplementary Table 4**) denoting CRC stem cell populations, largely compiled by Qin et al.^51^, GSVA was performed with pseudobulked CRC metacohort tumor samples and expression scores for each signature used for correlation analysis. Results yielded two phenotypic groups based on significant correlations between signatures, one of which aligned with the ASCL2^+^ transcriptome (**Fig. 3E**). These findings indicate the ASCL2^+^ phenotype shares similarities with previously defined progenitor cells, MEX3a^+^ reverse ISCs, LGR5^+^/MEX3a^+^ DTPs, proCSCs, and SSC1/2a proliferating ISC-like cells, among others^52–55^. Additionally, based on negative correlations the ASCL2^+^ phenotype is noted to be distinct from that seen in latent CLU^+^ CBCs, YAP induced fetal programed cells, revCSC, and metastasis-initiating EpiHR cells^51,56^. As implied by several signatures, comparison of stratified cell populations in the CRC metacohort confirmed ASCL2^+^ cells are a highly proliferative cell population (**Supplementary Fig. 2C**). Together the transcriptomic characterization revealed ASCL2^+^ cells are phenotypically proliferative, high plasticity, stem-like cells, with high WNT and low MAPK/YAP signaling, and represents a cell state present in a number of stem cell phenotypes previously identified in CRC.

To functionally confirm our transcriptomic findings of ASCL2^+^ cell features, we established a lentivirus based system to enable visualization and study of ASCL2^+^ cells. Given its superior efficacy, the STAR sequence from Oost et al.^36^ was utilized as the ASCL2 promoter to drive expression of mNeonGreen (‘GFP’ for simplicity) in ASCL2^+^ cells (**Fig. 4A**). CRC cell lines (B1003, WiDr, DLD1) were transduced with the designed lentivirus to establish ASCL2^GFP^ models. Cells from the generated models were flow-sorted to collect the GFP^+^ and GFP^-^ populations, and analysis by qPCR confirmed GFP^+^ cells had greater ASCL2 mRNA expression (**Supplementary Fig. 3A-B**). Although both groups displayed similar ASCL2 protein levels, GFP was significantly higher in the GFP^+^ cell group, proving the ASCL2^GFP^ system properly denotes cells with high ASCL2 activity (**Supplementary Fig. 3C**). With the system validated, GFP^High^ and GFP^Low^ cell populations were collected by flow-sorting to determine if ASCL2^+^ (GFP^High^) cells have higher expression of stem-related genes. Using a 16-gene panel for testing, results showed ASCL2^+^ cells had elevated SOX9, OCT4, KLF4, EMP1, and LRIG1 expression, while the differentiated cell marker CDX2 was less expressed than in the ASCL2^-^ population (**Fig. 4B**). These data demonstrate ASCL2^+^ cells have greater activation of stem programs, confirming our transcriptomic profiling findings. A hallmark of cancer is the phenotypic plasticity of tumor cells, a feature which has been reported for ASCL2^+^ cells in CRC^57,58^. ASCL2^+^ cells displayed higher transcriptomic entropy indicating a state of high plasticity, therefore we leveraged our models to test if the ASCL2^+^ state is a static or dynamic phenotype. Cells from B1003 ASCL2^GFP^ were sorted and monitored by live-cell imaging to track changes in cell populations. Initially the GFP^High^ and GFP^Low^ groups showed homogenous GFP^+^ or GFP^-^ cell populations, but numerous GFP^+^ cells were observed in the GFP^Low^ group by day 5, indicating the ASCL2^+^ phenotype is a dynamic state achievable despite prior cellular phenotype (**Fig. 4C**).

**Figure 4.**
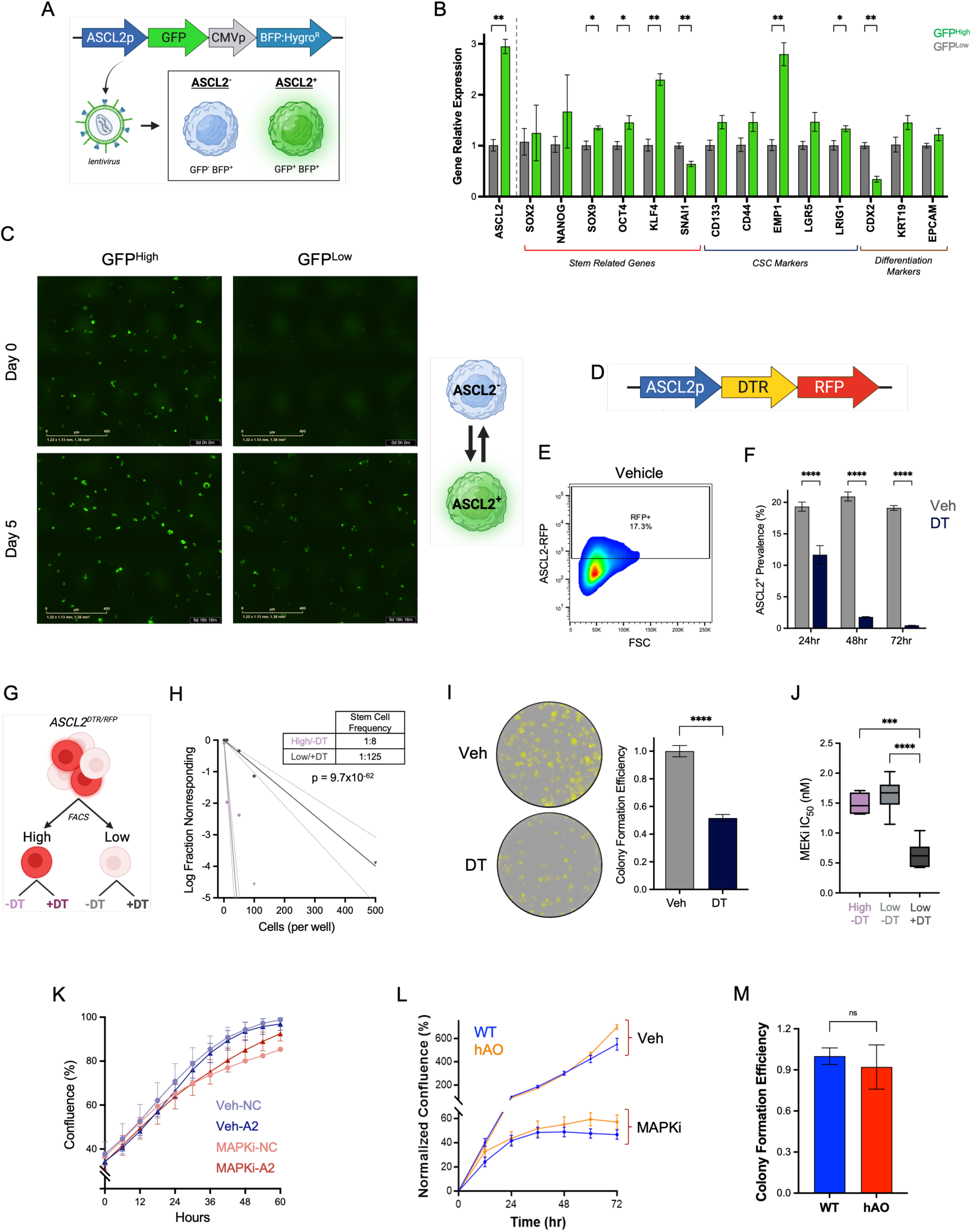
Functional characterization of ASCL2^+^ cells. **A)** ASCL2^GFP^ lentiviral vector, detailed in Supplementary Table 2, denotes ASCL2^+^ (GFP^+^/BFP^+^) and ASCL2^-^ (GFP^-^/BFP^+^) cells in CRC models. **B)** Gene expression of GFP^High^ (green) cells relative to GFP^Low^ (grey) cells from flow-sorted samples (n = 3) of WiDr ASCL2^GFP^ model; p < 0.05 (*), p ≤ 0.009 (**) (two-tailed t-test; mean ± SEM). **C)** Fluorescent cell imaging of B1003 ASCL2^GFP^ flow-sorted GFP^High^ and GFP^Low^ cells at Day 0 and Day 5; graphical illustration of the observed phenotypic plasticity. **D)** ASCL2^DTR/RFP^ lentiviral system for DT-ablation of ASCL2^+^ cells in CRC models. CT26 ASCL2^DTR/RFP^ model was used in following experiments (E-J). **E)** Representative flow plot of 72 hour vehicle treated CT26 sample used to quantify ASCL2^+^ cell population. **F)** Flow quantification of samples (n = 3) treated with vehicle (grey) or DT (black) for 24, 48, and 72 hours (two-way ANOVA with post-hoc Bonferroni; mean ± SEM; ****p.adj < 0.0001). **G)** RFP High (ASCL2^+^) and Low (ASCL2^-^) cells were collected by flow-sorting, and each population treated with vehicle or DT, forming four experimental groups. **H)** Spheroid forming limited dilution assay using sorted cells yielded stem-cell frequency of 1:8 cells for High/-DT (purple) group and 1:125 for Low/+DT (grey) group (chi-square ELDA test). **I)** Clonogenic assay of samples (n = 6) treated with vehicle (light grey) or DT (grey) to determine colony formation efficiency (two-sided t-test; mean ± SEM; p < 0.0001). **J)** MEKi IC_50_ (uM) of High/-DT (purple), Low/-DT (light grey), and Low/+DT (grey) groups (n = 6) from flow-sorted cells; all samples received 1 uM SOSi (BI-3406) in addition to MEKi (trametinib) treatment (one-way ANOVA with post-hoc Tukey; High/-DT vs Low/+DT p.adj = 0.0004, Low/-DT vs Low/+DT p.adj < 0.0001). Box plots depict median, first and third quartiles, and whiskers span from maximum to minimum value. **K)** Growth of B1003 cells treated with vehicle (blue) or MAPKi (red), and ASCL2 (A2) or noncoding (NC) siRNA. **L)** Relative growth of ASCL2 overexpressing (hAO; orange) or parental (WT; blue) B1003 cells treated with vehicle or MAPKi therapy for 72 hours. **M)** Colony formation efficiency by clonogenic assay of parental (WT) or ASCL2 overexpressed (hAO) B1003 cells (two-sided t-test; mean ± SEM).

The innate acquisition of the ASCL2^+^ state by CRC cells presents a major challenge for further functional testing, which requires maintenance of homogenous populations for comparison. To overcome the inherent plasticity, we generated a lentiviral ASCL2 specific diphtheria toxin receptor (DTR)^59,60^ and tdTomato (RFP) expression system, which enabled visualization and diphtheria toxin (DT) mediated cell ablation of ASCL2^+^ cells (**Fig. 4D, Supplementary Fig. 3D-E**). CT26 and MC38 were transduced with the ASCL2^DTR/RFP^ system and tested to ensure proper function. Flow cytometry of treated ASCL2^DTR/RFP^ cells showed successful ablation of ASCL2^+^ (RFP^+^) cells within 24 hours of DT treatment, and 93% depletion after 48 hours (**Fig. 4E-F**). Additionally, quantification of apoptotic RFP^+^ cells by live-cell imaging demonstrated a 95% ablation efficacy of ASCL2^+^ cells with DT-treatment (**Supplementary Fig. 3F**). The high efficiency achieved by DT treatment validates use of the ASCL2^DTR/RFP^ model to maintenance a homogenous ASCL2^-^ populations, through the continuous ablation of new ASCL2^+^ cells, enabling further characterization studies. A key feature of ASCL2^+^ cells is their stem-like phenotype, as previously reported and suggested by our transcriptomic analysis. Therefore, to inspect the stem capacity of ASCL2^+^ cells, a spheroid forming limited dilution assay^61^ was performed using RFP^High^ (High) and RFP^Low^ (Low) cells from the CT26 ASCL2^DTR/RFP^ model treated with vehicle or DT (**Fig. 4G, Supplementary Fig. 3E, 4A**). Resulting stem cell frequency was significantly higher for the High/-DT group (ASCL2^+^ population) than the Low/+DT (ASCL2^-^ population) group (p = 9.7x10^-62^), demonstrating ASCL2^+^ cells have greater stem cell capacity (**Fig. 4H**); similar findings were obtained when unsorted cells were treated with DT or vehicle (p = 1.2x10^-81^). It should be noted that stem cell frequency for the Low/-DT group, which represents an ASCL2^-^ population with intact ability to generate ASCL2^+^ cells, was not significantly different (p = 0.11) from the High/- DT group (**Supplementary Fig. 4B**). In a WiDr ASCL2^GFP^ model, no meaningful difference in stem cell frequency was observed between GFP^High^ and GFP^Low^ groups (**Supplementary Fig. 4C**). These findings emphasize that without mitigating the inherent ASCL2 phenotypic plasticity, characteristics unique to the ASCL2^+^ phenotype cannot be properly isolated. We also used colony formation assays as a second method to assess stem-like ability and further validate the ASCL2^+^ phenotype. As expected, assayed CT26 ASCL2^DTR/RFP^ cells showed that elimination of ASCL2^+^ cells by DT-treatment significantly reduced colony formation efficiency (**Fig. 4I**), supporting our overall findings. Given the association with worse clinical outcomes and the identified similarities with DTP cell types, we next investigated the tolerability of the ASCL2^+^ phenotype to MAPKi therapy. To assess MAPKi therapy response, flow sorted ASCL2^DTR/RFP^ cells were used to compare cell viability between groups (**Fig. 4G**). Both the High/-DT and Low/-DT groups displayed similar response to MAPKi (p = 0.61), but the Low/+DT group showed lower MEKi IC_50_ tolerability, indicating higher MAPKi sensitivity in ASCL2^-^ cells compared to ASCL2^+^ (p = 0.0004) or mixed (p < 0.0001) cell populations (**Fig. 4J; Supplementary Fig. 4D-E**). These findings reveal cells with the ASCL2^+^ phenotype have increased MAPKi therapy resistance, and sensitivity of ASCL2^-^ populations is neutralized by emergence of new ASCL2^+^ cells, highlighting the negative impact of plasticity on treatment response.

With the transcriptomic and functional characterization, we demonstrate that the ASCL2^+^ phenotype is a dynamic state and denotes a proliferative, stem-like population with high plasticity, and inherent MAPKi resistance, prompting us to define these cells as a distinct Adaptive Plasticity Tumor (APT) state and refer to them hereafter as APT cells. Lastly, we tested if ASCL2 produces the observed phenotype or is only a marker of APT cells. APT cells have higher MAPKi tolerability, however ASCL2 siRNA knockdown showed no impact on the response of CRC cells to MAPKi therapy (**Fig. 4K**). Similarly, overexpression of the human ASCL2 gene (hAO) did not impact stem-like ability (colony formation efficacy) or MAPKi therapy response of cells (**Fig. 4L-M, Supplementary Fig. 4F**). As such, these findings suggest the APT state is not primarily driven by ASCL2 alone, but ASCL2 is part of the phenotypic program and functions as an APT cell marker.

### MAPKi therapy enriches APT cells independent of CRC driver mutation

The effect of MAPKi therapy on the innately resistant APT cells was explored using CRC ASCL2^GFP^ models and monitoring for cell population changes. As such, B1003 ASCL2^GFP^ samples treated with MAPKi (BRAFi + EGFRi) or vehicle were processed by flow cytometry to quantify the APT (GFP^+^) cells (**Fig. 5A**). Results showed MAPKi therapy produced a significant enrichment of APT cells after 48 hours of treatment relative to the control group (**Fig. 5B**). Additional testing of ASCL2^GFP^ human (DLD1, WiDr) and murine (CT26) models showed that baseline APT cell prevalence varied across models, and each demonstrated significant increase in APT cell density with MAPKi therapy (**Fig. 5C**). These findings indicate MAPKi therapy enrichment of APT cells, a high-risk cell population, occurs regardless of driver mutation or treatment regimen, and represents a widespread process impacting CRC.

**Figure 5.**
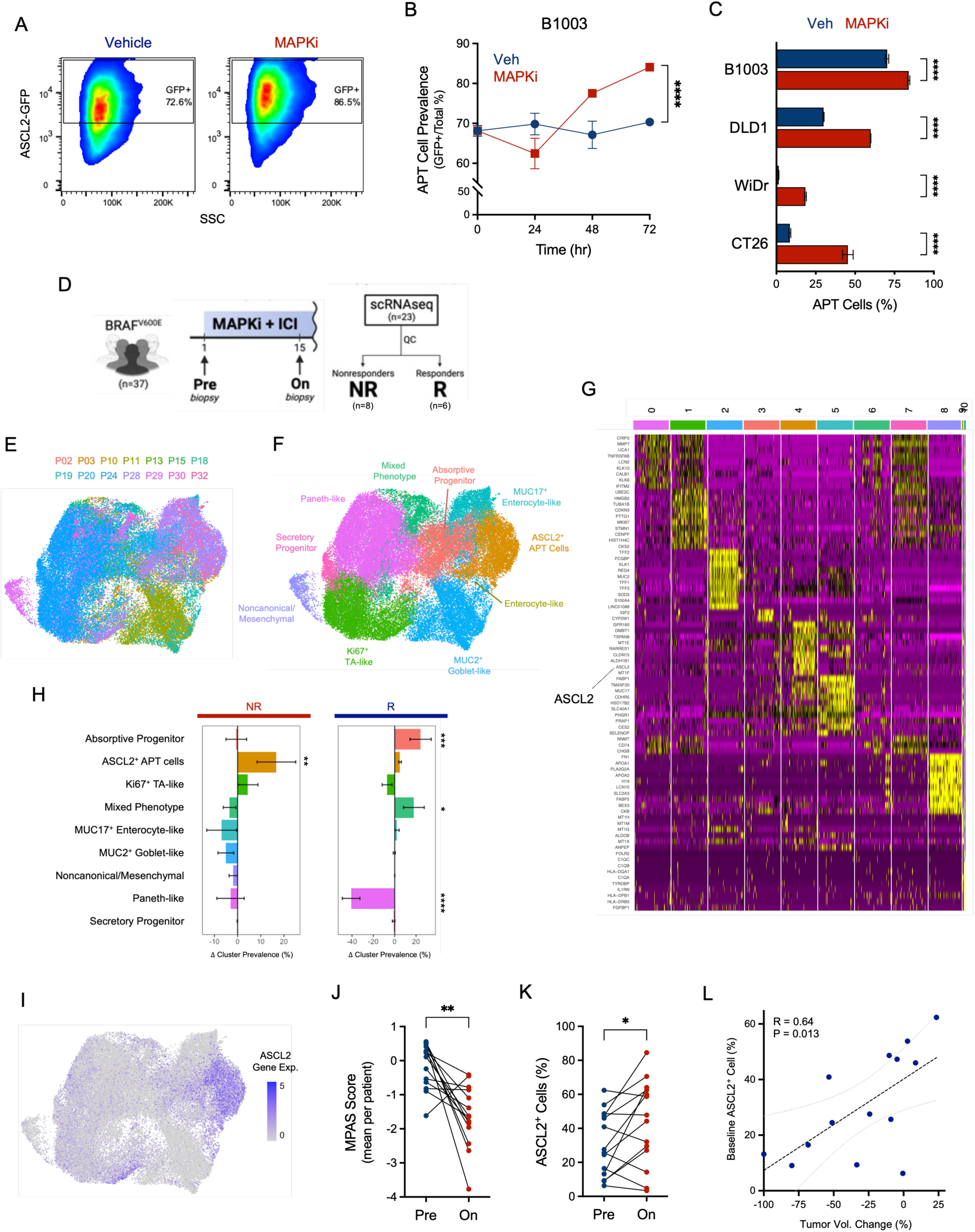
APT cells are enriched by MAPKi therapy. **A-B)** Flow cytometry (**A**) of B1003 ASCL2^GFP^ cells treated with vehicle or MAPKi (encorafenib, cetuximab), and samples (n = 3) collected at different timepoints to quantify APT (GFP^+^) cell population. **B)** Resulting prevalence of APT cells in vehicle (blue) and MAPKi (red) groups shown for each timepoint (two-sided t-test; mean ± SD). **C)** APT cell prevalence of CRC models treated for 72 hours with vehicle (blue) or corresponding MAPKi (red) therapy (n = 3, two-sided t-test; mean ± SD). **D)** BRAFi patient cohort study design (NCT03668431) in which pre- and on-treatment samples were collected for patients; 14 paired samples remained after processing scRNAseq data from 8 nonresponders (NR) and 6 responders (R). **E-F)** Batch corrected UMAP of scRNAseq tumor cells colored by patient (**E**) or cell cluster (**F**). **G)** Top 15 DEGs for each cluster identified by unsupervised analysis, with significant upregulation (yellow) of ASCL2 seen in cluster 4 cells. **H)** Average change in cluster prevalence between treatments (OnTx – PreTx) with patients grouped by response; clusters with significant changes are shown p.adj < 0.05 (*), p.adj < 0.005 (**), p.adj < 0.0005 (***), p.adj < 0.0001 (****) (two-way RM ANOVA with post-hoc Bonferroni; mean ± SE). **I)** UMAP of normalized ASCL2 expression in patient (n = 14) tumor cells. **J)** Average MAPK signaling (MPAS score) of tumor cells per patient at baseline (pre) and on MAPKi therapy (two-sided paired t-test; p = 0.0017). **K)** Cells classified by ASCL2 expression and percent ASCL2^+^ cells determined per sample; ASCL2^+^ cell prevalence compared by patient between treatments (two-sided paired t-test; p = 0.034). **L)** Pearson correlation (two-sided) of baseline (Pre) ASCL2^+^ cell prevalence and tumor volume change for each patient; simple linear regression trendline (black) and 95% CI (grey) are shown.

Clinical evidence of MAPKi driven APT cell enrichment was investigated using the scRNAseq data from the BRAFi patient cohort. After data processing 14 pairs of pre-treatment (PreTx) and on-treatment (OnTx) samples remained from 8 nonresponders (NR) and 6 responders (R), as determined by their clinical response (**Fig. 5D-E, Supplementary Fig. 5A**). Unsupervised clustering of tumor cells yielded 11 distinct clusters that were characterized by DEG analysis to identify their corresponding phenotypes (**Fig. 5F, Supplementary Fig. 5B-E**). Interestingly, ASCL2 was one of the top upregulated genes for cluster 4, a cluster with an ISC-like or early progenitor cell phenotype, which represented the ASCL2^+^ APT cell population (**Fig. 5G**). Tumor cell composition varied between patients (**Supplementary Fig. 5F**) and MAPKi treatment increased ASCL2 expression in the APT cell cluster (**Supplementary Fig. 6B-C**). Changes in tumor composition with MAPKi treatment was assessed by computing normalized fold-change (OnTx vs. PreTx) values of each identified cell phenotype for each patient. In both the NR patients (n = 8) and combined patients (n = 14), the ASCL2^+^ APT cell cluster was significantly enriched in tumors after initiating MAPKi therapy, whereas R patients had enrichment of absorptive progenitor and mixed phenotype cell clusters (**Fig. 5H, Supplementary Fig. 6D**). To further validate, a second approach was used where cells were classified based on ASCL2 expression as APT or nonAPT cells, and APT cell prevalence determined for each tumor sample (**Fig. 5I**). Paired comparison by patient demonstrated higher APT cell frequency in OnTx samples relative to PreTx, confirming that MAPKi therapy enrichment of APT cells is indeed observed clinically (**Fig. 5J**). Further analysis using clinical outcomes revealed that higher APT cell prevalence before treatment correlated with lower tumor volume change (r = 0.64, p = 0.013), indicating patients with higher baseline APT cell density were less responsive to MAPKi therapy (**Fig. 5K**). Of note, MAPKi therapy did not enrich LGR5^+^ or LRIG1^+^ cells in patient tumors, and neither baseline cell prevalence were predictive of therapy response (**Supplementary Fig. 6E-H**). Overall, we provide preclinical and clinical data that MAPKi-based therapy results in the enrichment of a resistant APT cell population, and show that baseline APT cell density predicts MAPKi therapeutic response in patients.

### APT phenotype is induced in CRC cells by MAPKi treatment

Due to the phenotypic plasticity of CRC cells, MAPKi therapy enrichment of APT cells could be the product of a selective or inductive process. To help elucidate the responsible process, flow sorted B1003 ASCL2^GFP^ APT cells (GFP^High^) and regular GFP^Low^ cells were treated with MAPKi and the resulting cell population re-analyzed by cytometry. Although MAPKi therapy did not alter the APT (GFP^+^) cell density in the plated APT (GFP^High^) cell samples, the sorted GFP^Low^ cell samples showed a large increase in the APT (GFP^+^) cell population with MAPKi treatment (**Supplementary Fig. 7A**). These findings indicated MAPKi treatment shifts basal APT cell equilibrium, resulting in cell enrichment by inducing the APT state in CRC cells. As such, we developed a single cell BarcodeSeq (scBarcodeSeq) platform to further investigate if MAPKi treatment enriches APT cells through phenotype induction. By combining molecular barcoding and single cell sequencing, scBarcodeSeq enables tracing individual cells across samples and comparing their transcriptomic profiles to gain insight on the impact of treatment at the single cell level (**Fig. 6A**). A barcoded CRC PDX model with a BRAF^V600E^ mutation, B1003b, was generated (outlined in methods) and mice with subcutaneous tumors were treated with MAPKi (encorafenib, cetuximab) or vehicle, and samples submitted for 10x Chromium at end-of-study for scBarcodeSeq (**Fig. 6B**). After processing the samples and sequencing data to incorporate cell barcodes, the resulting B1003b scBarcodeSeq data consisted of 10,952 cells with 6,985 unique barcodes (**Fig. 6C**).

**Figure 6.**
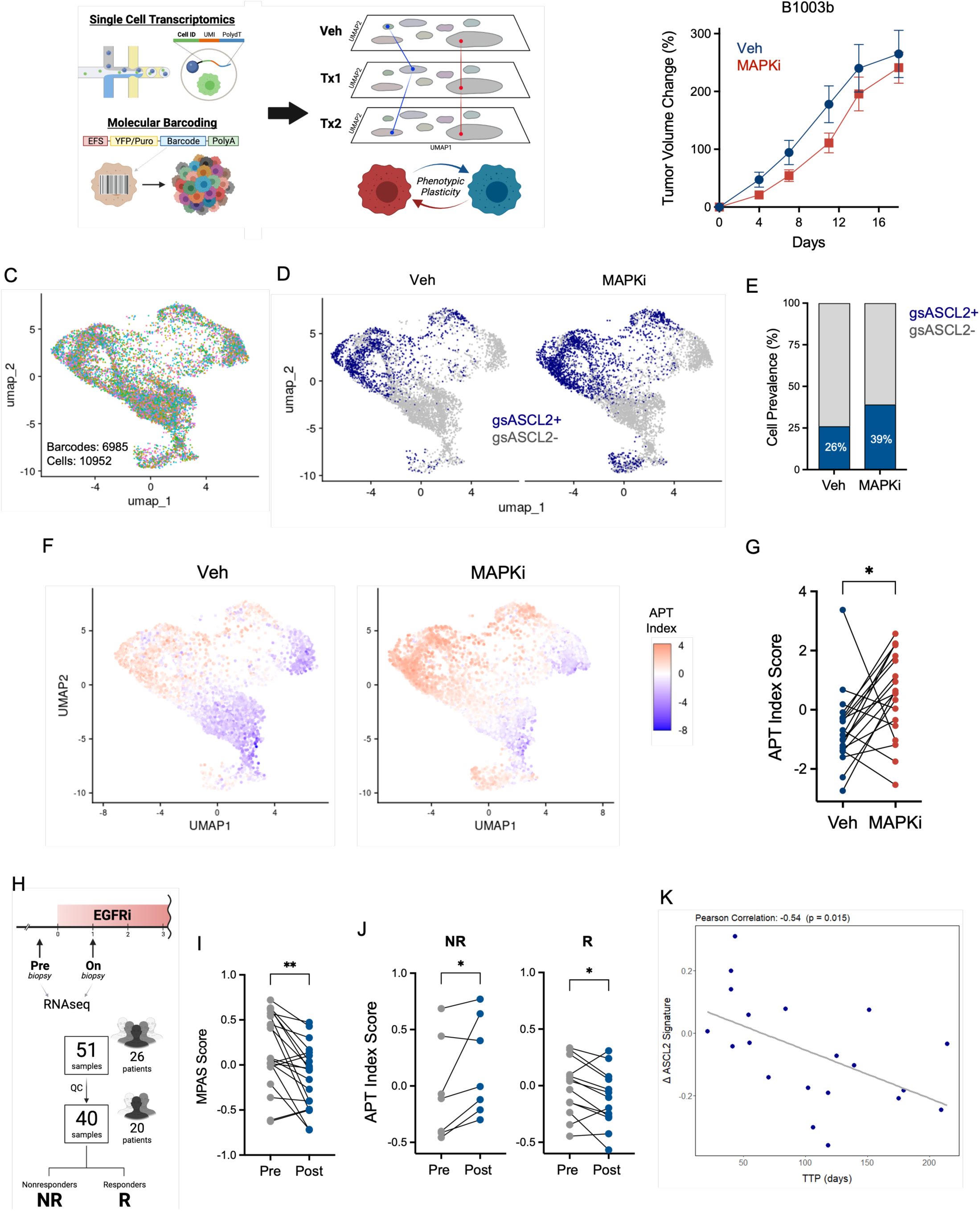
MAPKi therapy induces APT phenotype in CRC cells. **A)** Single cell BarcodeSeq (scBarcodeSeq) uses molecular barcoded tumors and single cell transcriptomics to enable comparison of individual cells across samples. **B)** Barcoded B1003 (B1003b) tumor volume change in mice (n = 5) treated with vehicle (blue) or MAPKi (encorafenib, cetuximab) therapy (red) for 21 days (mean ± SEM). **C-E)** UMAP from scBarcodeSeq of B1003b samples colored by barcode ID (**C**), consisting of 10,952 tumor cells with 6,985 unique barcodes. **D)** Tumor cells were classified by gsASCL2 expression as positive (blue) or negative (grey) for vehicle and MAPKi treated samples. **E)** Prevalence of cell types shows higher ASCL2^+^ cell density with MAPKi (39%) compared to vehicle (26%) treatment (chi-square, p < 0.0001). **F-G)** APT Index signature scores for vehicle and MAPKi treated cells, shown for individual cells using UMAP (**F**), and (**G**) averaged by cell with matching barcodes in each treatment group (Wilcoxon matched pairs signed rank test; p = 0.018). **H)** EGFRi rechallenge cohort of RNAseq samples from patients before (Pre) and after (Post) starting treatment; after QC, paired data remained for 7 nonresponders (NR) and 13 responders (R). **I)** MAPK activity score (MPAS) expression in patients (n = 20) for pre (grey) and post (blue) treatment samples (Wilcoxon matched-pair signed rank test; p = 0.001). **J)** APT (ASCL2) Index expression at each timepoint for NR (p = 0.031) and R (p = 0.048) patients (Wilcoxon matched-pair signed rank test). **K)** Pearson correlation (two-sided) of delta APT Index (Post - Pre), representing extent of ASCL2 enrichment, and Time-to-Progression (TTP) per patient (n = 20); simple linear regression trendline (grey).

Analysis of the B1003b scBarcodeSeq data where cells were classified by gsASCL2 expression to denote APT cells (ASCL2^+^) showed MAPKi therapy enriched the APT cell population (39% vs. 26%; p < 0.0001), as expected (**Fig. 6D-E**). Cells with matching barcodes in the vehicle and MAPKi-treated samples were compared by barcode ID to assess changes in gene expression. Despite the modest *in vivo* tumor response to MAPKi therapy, evaluation of MAPK signaling using MPAS (MAPK Pathway Activity Signature) score revealed large reduction of MAPK activity in treated cells relative to corresponding control cells, indicating successful therapeutic pathway suppression (**Supplementary Fig. 7B-C**). The APT Index was used to gauge expression of the APT cell phenotype in matching barcoded cells, and analysis showed MAPKi treatment produced a significant increase in the expression of APT Index in matched cells (**Fig. 6F-G**). Together, these findings indicate that inhibition of MAPK activity resulted in the induction of the APT cell phenotype, producing the observed enrichment in the patient-derived CRC tumors.

MAPKi therapy enriched APT cells in the BRAFi patient scRNAseq cohort (**Fig. 5J**), and GSEA showed that treatment in nonresponders produced increased tumor expression of the APT phenotype (gsASCL2, NES = 1.25, p.adj = 3.69 x10^-5^) compared to responder patients (**Supplementary Fig. 6I**). To further gauge the clinical relevance of these findings we utilized RNAseq data from a set of clinical trials (NCT03087071, NCT04616183) where paired tumor samples were collected from patients (n = 26) with BRAF/KRAS^WT^ mCRC prior to initiating EGFRi therapy rechallenge (Pre) and after one month (Post) of treatment (**Fig. 6H**). Of the 51 collected samples, 20 data pairs remained after data processing, corresponding to 7 nonresponder (NR) and 13 responder (R) patients. Expression of gene signatures in each sample was computed by GSVA to assess transcriptomic changes across patient-matched tumors. Comparison of MAPK signaling (MPAS signature) between treatment samples demonstrated significant reduction of MAPK activity, and successful therapeutic effect (**Fig. 6I**). APT phenotype expression was determined using the APT Index (ASCL2 Index) signature score, showing a significant increase in APT Index in NR patients, while R patients displayed a modest reduction (**Fig. 6J**). Of import, the change in tumor APT signature expression (Δ APT signature) was inversely correlated with time-to-progression (TTP), indicating greater MAPKi therapy APT phenotype enrichment resulted in less durable responses (**Fig. 6K**). Although not at the single cell level, this data provides insights into the transcriptomic profile of cells in CRC tumors, showing EGFRi therapy effectively suppressed MAPKi signaling but resulted in elevated expression of APT cell program in the NR patients. Results of GSEA similarly demonstrated MAPKi treatment enriched APT phenotype expression in NR compared to R patients (**Supplementary Fig. 5J**). Additionally, a seperate RNAseq dataset of patients (n = 5) with KRAS^G12C^ mCRC, with samples collected before receiving MAPKi treatment (baseline) and at the time of disease progression, showed enrichment of APT phenotype in treated tumors at progression (**Supplementary Fig. 5K**). Altogether, these findings provide further support that therapeutic MAPK pathway inhibition induced APT cell program expression in a patient subset who had less durable response.

### APT cells reduce MAPKi response durability in CRC

Leveraging our ASCL2^DTR/RFP^ models, we explored the impact of APT cells on MAPKi therapy by depleting the APT cell population. Since MAPKi treatment caused enrichment of APT cells, we first tested if DT-ablation was able to sufficiently deplete APT cells in samples concurrently treated with MAPKi. Live-cell imaging of CT26 ASCL2^DTR/RFP^ cells stained with a fluorescent apoptosis dye showed that DT-treatment achieved 75-90% ablation efficacy of APT cells when samples were co-treated with MAPKi therapy (**Fig. 7A**). To confirm these findings, quantification of APT cell (RFP^+^) prevalence was performed by flow cytometry in treated samples. As expected, MAPKi treatment produced an enrichment in APT cells, however co-treatment with DT significantly reduced the APT population, ablating approximately 80% of cells (**Fig. 7B-C**). As such, the DT-ablation system successfully depleted APT cells in MAPKi co-treated samples, indicating it can be used to investigate the effect APT cells have on MAPKi therapy. As the DTR system can only be used in murine lines, to test a second CRC model of different mutation background than CT26 (BRAF/KRAS^WT^), we obtained the syngeneic ABPS (APC^-^, BRAF^V600E^, TP53^fl/fl^, SMAD4^-^) CRC cell line^16^ for additional study. Due to native tdTomato expression of ABPS, a new DT-APT cell ablation system was generated, switching the fluorescent protein to mNeonGreen (GFP), and used to establish an ABPS ASCL2^DTR/GFP^ model. For MAPKi treatment, ABPS cells received encorafenib (BRAFi), trametinib (MEKi) and cetuximab (EGFRi). Cell viability and growth were monitored to determine if APT cell ablation (+DT) altered the response to MAPKi therapy. Although APT cell depletion (DT treatment) alone resulted in a mild reduction in cell viability (**Fig. 7D, F**), cell growth was minimally impacted, displaying a slight delay in growth (**Fig. 7E, G**). In both models, combination of MAPKi therapy with DT-ablation produced a greater reduction in cell viability than MAPKi therapy, indicating depletion of APT cells improved MAPKi response (**Fig. 7D, F; Supplementary Fig. 8B-C**). Additionally, dual MAPKi and DT treatment displayed longer cell growth inhibition (**Fig. 7E, G**), as highlighted on day 10 where MAPKi samples were confluent and contained multiple APT (RFP^+^) cells, while MAPKi/DT samples remained <50% confluent and had no viable APT cells present (**Fig. 7H, Supplementary Fig. 8A**). Together, these data demonstrate APT cell depletion improved efficacy and prolonged response of MAPKi therapy, confirming that APT cells negatively impact MAPKi response in CRC.

**Figure 7.**
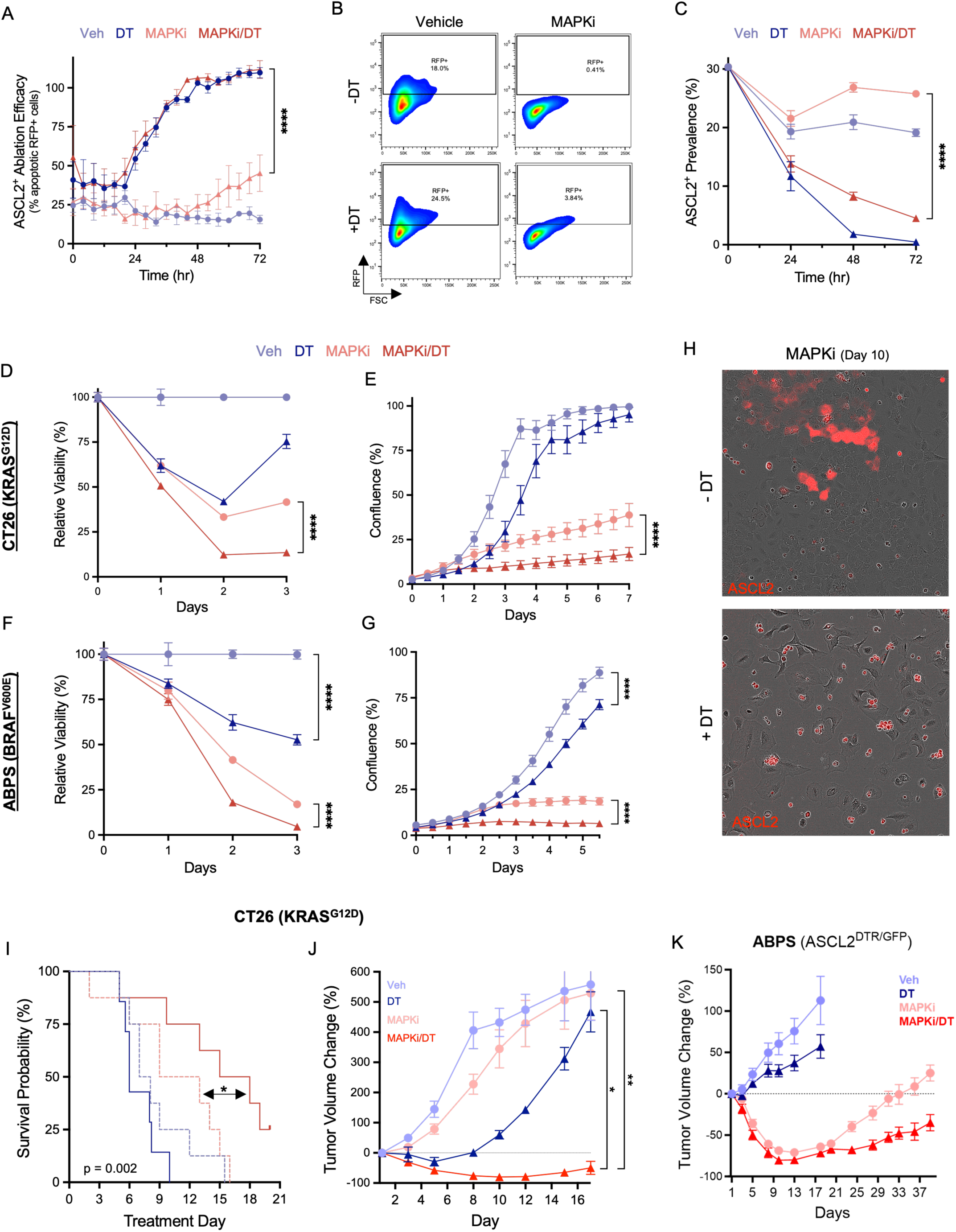
APT cell depletion improves MAPKi therapy response. **A)** APT (ASCL2^+^) cell ablation efficacy of CT26 ASCL2^DTR/RFP^ model treated with vehicle, DT, MAPKi (BI-3406, trametinib), or MAPKi/DT; cells stained with apoptosis (GFP) marker and samples (n = 6) monitored by IF cell imaging (AUC calculated per group then one-way ANOVA with post-hoc Tukey; mean ± SEM). **B-C)** Flow cytometry of CT26 ASCL2^DTR/RFP^ cells treated with vehicle or MAPKi, with (+DT) and without (-DT) concurrent APT cell ablation, and collected at 0, 24, 48, and 72 hours (**B**) to quantify APT (RFP^+^) cell population per sample (n = 3). **C)** Resulting APT cell prevalence in vehicle, DT, MAPKi, and MAPKi/DT groups at each timepoint (two-way ANOVA with Bonferroni; mean ± SD). **D-G)** For all plots, the MAPKi treatment group is shown in red and vehicle group in blue. Data from samples treated with DT are triangles and vehicle (-DT) are circles. **D, F)** Cell viability relative to vehicle for CT26 ASCL2^DTR/RFP^ (**D, E**) and ABPS ASCL2^DTR/GFP^ (**F, G**) samples (n = 6) treated with vehicle, DT, MAPKi, or MAPKi/DT (two-way ANOVA with Bonferroni; mean ± SD; ****p.adj < 0.0001). **E, G)** Similarly, both models were treated and cell growth monitored by well confluency (n = 10) for 1 week (one-way ANOVA of AUC with Tukey; mean ± SEM). **H)** Representative images from day 10 of ASCL2^DTR/RFP^ cells treated with MAPKi +/- DT, and monitored by fluorescent and brightfield imaging. **I)** Kaplan-Meier plot of survival in mice (n = 5) with CT26 ASCL2^DTR/RFP^ tumors treated with vehicle/-DT (mean = 8 days), vehicle/+DT (7 days), MAPKi/-DT (9 days), or MAPKi/DT (15 days); overall log-rank p = 0.002, and pairwise logrank comparison of MAPKi/+DT vs. MAPKi/-DT p.adj = 0.04. **J)** Tumor volume relative to day 0 for mice with CT26 ASCL2^DTR/GFP^ tumor volume change, relative to day 0, in mice (n = 8) treated with vehicle/-DT, vehicle/+DT, MAPKi/-DT, and MAPKi/+DT (Kruskal-Wallis test with Dunn’s multiple comparison; mean ± SEM; *p.adj = 0.024, **p.adj = 0.0047). **K)** Same as (**J**) but using ABPS ASCL2^DTR/GFP^ mice for study.

Treatment with MAPKi agents in mCRC results in the enrichment of APT cells, a unique cell population with increased resistance, which we found dampens response to MAPKi therapy. These findings suggest APT cells may contribute to the limited durability of MAPKi therapy that is observed clinically in CRC. As such, we conducted *in vivo* studies to determine if elimination of the APT cell population in CRC tumors alters the duration of response to MAPKi therapy in mice. Athymic nude mice with subcutaneous CT26 ASCL2^DTR/RFP^ tumors were treated with vehicle, DT, MAPKi (BI-3406, trametinib), or with the combination (MAPKi/DT) to assess impact on survival and tumor growth. In CT26, MAPKi treatment had minimal effect on tumor volume compared to control, while ablating APT cells with DT decreased tumor volume and resulted in delayed tumor growth. Despite the modest response obtained by MAPKi, combined MAPKi/DT treatment showed a significant reduction in tumor volume and longer interval of suppressed tumor growth (**Fig. 7J**). A separate study with similar results, showed median survival of mice treated with vehicle or DT was 6-8 days, while MAPKi treatment provided a small benefit (11 days). Interestingly, MAPKi/DT treatment significantly extended survival (17 days) in mice relative to MAPKi therapy alone (p = 0.04), indicating depletion of APT cells in CRC tumors improves MAPKi response (**Fig. 7I**). Similarly, mice with ABPS ASCL2^DTR/GFP^ subcutaneous tumors received vehicle, DT, MAPKi (encorafenib, trametinib, cetuximab), or dual MAPKi/DT treatment. Unlike CT26, ABPS tumors readily responded to MAPKi therapy, and DT treatment produced a small tumor volume decrease. However, ablation of APT cells in MAPKi treated (MAPKi/DT) tumors resulted in more durable tumor growth suppression and successfully prolonged response to MAPKi (**Fig. 7K**). Combined, these studies confirm that APT cells mitigate MAPKi therapy response in CRC and highlight their important role in tumor biology. Altogether, we demonstrate MAPKi therapy activates intrinsic cell programs inducing APT cell phenotype, which becomes enriched in tumors, due to their increased resistance, ultimately limiting durable response to MAPKi therapy (**Supplementary Fig. 8D**).

## Discussion

MAPK targeting therapies have become a key component in the treatment of patients with mCRC, but use continues to be limited by short-lived therapeutic response^1,24^. Multiple studies have reported acquired resistance through subclonal *de novo* mutations in a subset of mCRC patients, although the cause remains uncertain^21,23^. Here, we identify new tumor biology and present novel insights of the impact tumor plasticity has on MAPKi therapy response durability in CRC. Our findings elucidate a novel process where MAPKi therapy activates innate plasticity of CRC cells, prompting dedifferentiation into an APT state, a distinct phenotype with increased resistance to MAPKi, which becomes enriched in tumors, decreasing the therapeutic effect and shortening response durability of MAPKi therapy.

Analysis of clinical trial data^16^ (NCT03668431) from patients with BRAF^V600E^ mCRC showed transcriptional activation of stem-related programs and increased ASCL2 expression with MAPKi treatment. Similar findings were observed in patient-derived CRC models, and additionally showed higher ASCL2 activity, a surrogate marker of the APT state, in treated tumors. ASCL2, a transcription factor, is known to serve important functions in several organ systems^62,63^, and its association with worse outcomes in mCRC implies ASCL2 may play an important role in CRC. ASCL2 has been shown to alter the tumor microenvironment in CRC, and ASCL2 expressing cells reported to have stem-like, neuroendocrine, and reserve ISC features, however a detailed description of ASCL2 cells has not been established^42,45,48,64–66^. Given our findings, we explored the corresponding phenotype using the specialized ASCL2 reporter, generated by Oost et al.^67^, to identify ASCL2^+^ cells. Phenotype characterization revealed a proliferative cell with high plasticity and stem capacity, representing a dynamic state achieved by innate plasticity. As such, the distinct phenotype was renamed Adaptive Plasticity Tumor (APT) cells, and were found to have increased resistance to MAPKi therapy. The APT phenotype shared transcriptomic similarities with other reported CRC stem-like cell phenotypes highlighted its biological relevance. Additionally, correlations with LGR5, progenitor and MEX3A signatures indicate the APT phenotype represents an activated reserve population of tumor ISC-like cell in CRC, which is consistent with the APT cell characteristics we observed. Although in our results ASCL2 does not appear to independently drive the APT phenotype, further studies are needed to determine the transcriptional drivers and confirm if ASCL2 is only a biomarker. Nonetheless, clinical evidence of stem program activation in MAPKi treated tumors led to the identification of APT cells and elucidating their unique characteristics, which suggested APT cells may impact patient outcomes.

Features of the APT phenotype suggest increased ASCL2 expression resulting from MAPKi treatment may be part of an adaptive response program to alter cell state and minimize impact of therapy. Leveraging lineage tracing tools and NGS technologies, scBarcodeSeq enabled direct comparison of matching barcoded tumor cells across treatments to assess transcriptomic changes in individual cells. Using this approach, we find MAPK pathway inhibition in cells resulted in higher expression of the APT phenotype, and, with additional findings, conclude that MAPKi therapy induces the APT cell state. Additionally, MAPKi treatment caused enrichment of the APT cell population in multiple human and murine CRC models. Our findings indicate that CRC treatment with MAPKi enriches APT cells primarily through a Lamarckian model of phenotype induction. Although a Darwinian-like selection may also contribute to some extent, these results highlight the critical role tumor plasticity plays in CRC tumor biology. Importantly, evidence of APT cell enrichment by MAPKi treatment was observed in three independent sets of patients with mCRC – the BRAFi patient cohort (BRAF^V600E^), the EGFRi rechallenge cohort (KRAS/BRAF^WT^), and the G12Ci progression cohort (KRAS^G12C^). Combined with our preclinical findings, these results confirm MAPKi enrichment of APT cells occurs irrespective of driver mutations or treatment regimen, indicating broad impact in mCRC patients and high clinical relevance. Further, clinical data showed pre-treatment APT cell prevalence is predictive of tumor response, and greater APT enrichment is associated with less durable response. As such, future prospective trials would be beneficial to investigate APT cell density as a potential biomarker of MAPKi therapy response.

Despite the common use of MAPKi therapy in the treatment of mCRC, few studies have explored its effect on stem cells and tumor plasticity^68–70^. Chemotherapy has been widely reported to enrich various stem cell populations in CRC, and was recently shown to impact metastases by increasing fetal progenitor state-derived noncanonical phenotypes, both of which are associated with worse outcomes^35,71–73^. Although similar investigations using targeted therapies have been limited, one study demonstrated targeted therapy alters expression of two identified colonic stem cell phenotypes^51^. As such, our findings help define an important tumor cell population (APT) and provide valuable insight on the effects of MAPKi therapy on stemness and plasticity in CRC tumors. However, as MAPKi therapy is increasingly used in combination with chemotherapy, a remaining question that should be explored is the impact of chemotherapy and MAPKi plus chemotherapy on the APT cell populations in CRC.

Our data shows MAPKi therapy results in engagement of plasticity, shifting the phenotypic equilibrium of cells to enrich the population with decreased MAPKi sensitivity. Further, ablation of APT cells in models treated with MAPKi therapy showed greater efficiency than MAPKi alone, confirming the negative impact APT cells and tumor plasticity have on therapy response. Additionally, MAPKi therapy with concurrent APT cell depletion increased treatment efficacy, prolonged response durability, and improved outcomes in our preclinical studies. Of note, these results were observed across CRC models despite different mutation background (KRAS/BRAF^WT^, BRAF^V600E^) and MAPKi treatment, highlighting the ability of APT cells to mitigate MAPKi therapy response independent of tumor mutation status. Altogether, we report a new mechanism where therapeutic MAPK pathway inhibition activates cellular plasticity resulting in CRC cells entering an APT state, which becomes enriched in tumors due to their MAPKi resistance and reduces therapeutic response.

Despite the array of supportive data, there are limitations to this study. Foremost, there are few clinical studies with paired samples collected before and after patients with mCRC receive MAPKi therapy, and is further decreased by lack of data accessibility, severely limiting usable data for our analysis. Due to the BRAFi patient cohort receiving both MAPKi and immune checkpoint inhibitor, immunotherapy represents a potential confounding variable impacting our observations. The study was used given data scarcity, but chances our findings are nonspecific are greatly minimized since additional clinical and pre-clinical data only received MAPKi therapy. Use of the ASCL2^DTR^ system was limited to mouse CRC models due to the innate sensitivity of human cells to DT. Although an alternative system compatible with human models was tested, it failed to produce adequate ablation of APT cells. Therefore, improved response of MAPKi therapy upon APT (ASCL2^DTR^) cell depletion was confirmed in two murine models, but additional validation in human CRC models was not possible. Nonetheless, this work provides compelling findings outlining the detrimental effect tumor plasticity has on MAPKi therapy response in CRC.

Cancer cells can more readily adapt to different conditions than healthy cells, suggesting malignancy is accompanied by greater plasticity, although where along the tumorigenesis sequence this is obtained remains unknown^74–80^. Although advances are being made in understanding the progression of CRC, tumor plasticity remains a major challenge in oncology. Overall, these findings identify a dynamic cell state expressed by a tumor cell population that mitigates the response durability of MAPKi therapy in CRC. While the mechanism of tumor cell plasticity that links MAPK pathway inhibition and APT state induction, remains an unsolved question, this work paves the way to a better understanding of plasticity and targeting of this cell population to improve therapy response durability in patients.

## Data Availability

BRAFi clinical trial scRNAseq data was obtained from the Single Cell Portal (SCP2079). CRC metacohort was generated using data from Synapse (syn26720761, syn26844071) and our prior published MDACC dataset (GSE231559). ASCL2 gene signature was generated using GSE99133. Data from B1003b scRNAseq will be uploaded to GEO after publication of manuscript. Code script used for data processing and analysis can be provided upon request to corresponding author.

## Acknowledgements

We would like to acknowledge the contributions of Eduardo Vilar-Sanchez, George Calin, Jessica Bowser, and Michael Kim in the development of this project, in addition to the individuals at the Translational Research to AdvanCe Therapeutics and Innovation in Oncology (TRACTION) for their help and expertise. OEV was supported by the CPRIT Training Program (RP210028). This work was supported in part by the UTHealth Houston/MD Anderson Cancer Center Graduate School of Biomedical Sciences Kopchick Fellowship awards, and a CPRIT Single Cell Genomics User Group Grant 2022. SK and this work were supported by the Gastro-Intestinal Research Foundation (FP18308), Stand Up 2 Cancer (FP14745), Department of Defense (CA2308461), Gastrointestinal Cancer SPORE (P50CA221707), and NIH (R01CA262805). JPS was supported by the Col. Daniel Connelly Memorial Fund, the Andrew Sabin Family Fellowship Award, the Cancer Prevention & Research Institute of Texas as a CPRIT Scholar in cancer Research (RR180035, RP240392), and Conquer Cancer (2022CDA-7604125121). The Flow Cytometry and Cellular Imaging Core Facility was supported in part by the University of Texas MD Anderson Cancer Center and P30CA016672. The Advanced Technology Genomics Core was supported in part by the University of Texas MD Anderson Cancer Center, P30CA016672 and NIH 1S10OD024977-01. The Research Animal Support Facility was supported in part by the University of Texas MD Anderson Cancer Center and P30CA016672. The Single Cell Genomics Core Facility was supported in part by the facilities grant RP180684. BioRender used to generate several illustrations throughout manuscript.

## Methods

### BRAFi patient scRNAseq Data

Raw counts for epithelial cells and metadata files from the clinical trial (NCT03668431) were obtained from Single Cell Portal (SCP2079) and further processed with the R package Seurat v5^81,82^. Dataset consisted of samples collected prior to treatment, which were labeled “Pre”, and samples collected on day 15 of treatment, annotated “On”^16^. Quality control (QC) was performed by filtering out cells with mitochondrial gene content >25%, and samples with less than 5 cells were removed from the dataset, along with unpaired samples. Resulting dataset consisted of 55,928 cells from 14 paired patient samples. Data was normalized using SCTransform function, with mitochondrial genes fraction (percent.mt) regressed out, before Treatment and inter-patient batch effect was corrected using FastMNN data integration and a uniform manifold approximation and projection (UMAP) plot generated for data visualization. Metadata contained clinical outcomes of patients including progression free survival (PFS) and response status based on RECIST criteria, which was used to classify patients as responders (R) or nonresponders (NR). The processed data represented paired sampled from 8 nonresponders (NR) and 6 responders (R), and is referred to as the “BRAFi patient cohort”.

### BRAFi patient Pseudobulk Analysis

Due to high variability of total number of cells per sample additional cell filtering was performed to avoid large differences across samples in total reads after pseudobulking. All samples with < 50 cells were excluded, and patients were removed if one of the paired samples, either ‘Pre’ or ‘On’ had < 250 total cells. A total of 22 samples remained from 6 NR and 5 R patients. To minimize the significant batch effect observed in data processing, the SCT normalized counts were used for pseudobulking, in which cells were aggregated by sample and counts per gene summed to generate a pseudobulk count matrix. Differentially expressed gene (DEG) analysis was performed with DESeq2 accounting for patients, treatment and response (model: patient_n + response:patient_n + treatment*response)^83^. Principle component analysis (PCA) plot was generated using VST normalized data with DESeq2. DEG analysis was performed by contrasting NR (On vs. Pre) with R (On vs. Pre) to assess differences due to treatment between nonresponders and responders, and DEGs ranked using the Wald test statistic. Gene Set Enrichment Analysis (GSEA) was conducted using a curated list of 51 gene signatures reported to correlate with outcomes in CRC (**Supplementary Table 3**) and results were filtered using a false discovery rate (FDR) < 0.05 to identify top signatures^84^. Normalized expression of genes commonly used as CRC stem cell markers were obtained from DESeq2 and log scaled to assess change with treatment across all patients. Gene expression change was calculated per patient (On vs Pre) for each gene, without stratifying patients based on response. Statistical significance (p-value) for each gene of interest was extracted from the On vs Pre-treatment DEG analysis.

### BRAFi patient scRNAseq Analysis

Processed data from BRAFi patient cohort, outlined above, was further analyzed using Seurat v5. Unsupervised clustering of cells was performed using the FindClusters function with resolution set to 0.5 and produced 11 clusters and UMAP used for visualization. Differentially expressed genes for each cluster were identified with the FindAllMarkers function using a Wilcoxon rank sum test and the top 15 genes per cluster used to create a heatmap. To calculate the prevalence of cells in the clusters per patient, the number of cells in each Seurat cluster was determined for each sample and divided by the total number of cells in the sample. Fold change (FC) of cluster prevalence was computed per patient (On vs Pre) and log normalized, resulting log_2_FC values were shown by heatmap and averages computed per cluster for R and NR groups, as well as all patients combined. A two-way ANOVA with post-hoc Bonferroni was used to determine statistical significance; p-adjusted values of clusters with a significant change are displayed on heatmap. To evaluate MAPKi induced ASCL2 expression changes within cluster 4 cells, the scRNAseq data was pseudobulked by cluster, using the methods outlined above, and ASCL2 DESeq2 log normalized values for each sample compared using a two-tailed Mann-Whitney test. Analysis of APT cell prevalence change with treatment was performed as follows. Cells were classified based on their normalized expression of ASCL2 and denoted as ‘positive’ or ‘negative’. Percent of cells for each cell subtype was computed by sample, and ASCL2^+^ cell prevalence compared between timepoints by patient (n = 14) using a Wilcoxon matched-paired signed rank test. Baseline ASCL2^+^ cell prevalence for each patient was compared to the reported tumor volume change using a Pearson correlation test.

### CRC Metacohort scRNAseq Data

Paired tumor and normal adjacent tissue samples from patients with CRC from four independent patient cohorts were combined to form a metacohort of scRNAseq data. Consolidated normalized counts and corresponding metadata for three cohorts were published by Joanito et al., and data for the 51 combined patients was obtained from Synapse (syn26720761, syn26844071)^38^. Our group recently published a cohort of CRC scRNAseq data consisting of 9 patients with CRC (GSE231559)^34^, and epithelial cells from each dataset were merged and processed using Seurat. Cell QC filtering was performed to remove cells with mitochondrial gene percent > 50%, resulting in 59,727 cells. Cohort batch effect was corrected by normalizing counts using SCTransform and performing RPCA integration of layers to generate a batch corrected UMAP. Cells were classified based on scaled normalized gene expression of CRC stem cell markers, and annotated as ‘positive’ or ‘negative’ for each gene marker. Classified cells were divided based on tissue type (tumor, normal), and total number of cells for each stem marker phenotype were computed. Association of each stem marker with tumor cells was assessed by calculating odds ratio (OR), 95% confidence interval (CI), and a two-tailed Fisher exact test used for statistical significance. Coexpression of top 3 stem markers (ASCL2, LGR5, LRIG1) was assessed for tumor cells using the same cell annotations and determining the prevalence of each coexpressed phenotype.

### Generating ASCL2 Gene Signature

RNAseq data published by Oost et al. was obtained from GEO database (GSE99133)^36^. FASTQ files were aligned to human genome (GRCh38), annotated and counted using Rsubread package. Gene counts were analyzed with DESeq2 by comparing STAR^+^ (ASCL2^+^) and STAR^-^ (ASCL2^-^) samples to identify DEGs for the signature. Signature genes upregulated in ASCL2^+^/STAR^+^ samples, those with log2FC > 0, were labeled as the ASCL2^UP^ signature (gsASCL2, or gsASCL2-UP), whereas downregulated genes were used as a gsASCL2-DN signature. To calculate signature expression scores in scRNAseq data, the AddModuleScore function from

Seurat was used and resulting values for each gene signature were z-normalized by sample. Additionally, to more thoroughly capture the transcriptome of the ASCL2^+^ cell phenotype, an ‘ASCL2 Index’ was computed using the normalized gene signature scores (gsASCL2 – gsASCL2-DN). Validation of the ASCL2 gene signature was conducted using the CRC Metacohort tumor cell data, where gsASCL2 scores were found to significantly correlate with normalized ASCL2 gene expression (p = 0.008); similar findings were obtained with ASCL2 Index and using different datasets to confirm findings. After characterization of ASCL2^+^ cells, the ASCL2 Index was renamed the APT Cell Index.

### CRC Metacohort scRNAseq Analysis

Processed data from CRC Metacohort, outlined above, was further analyzed using Seurat as follows. Cells from normal tissue samples were removed to obtain a tumor cell dataset. Metacohort cells were classified by ASCL2 normalized gene expression and annotated as ‘positive’ or ‘negative’. Similarly, cells were annotated based ASCL2 gene signature expression using the AddModuleScore function and z-normalizing the gsASCL2 expression scores. Stem potency of cells in dataset was calculated using the CytoScore produced by the CytoTRACE2 package^85^. Cell entropy was computed with perCellEntropy function from the TSCAN package.

Scores of interest (gsASCL2, CytoScore, entropy) were averaged for ASCL2^+^ and ASCL2^-^ cells by sample, or using the gsASCL2 cell annotations, and difference in scores between cell phenotypes (ASCL2^+^ vs ASCL2^-^) assessed with a two-sided Mann-Whitney test. Correlations were tested by comparing averaged gsASCL2 and entropy scores per patient using a Pearson correlation.

For testing transcriptomic profile of ASCL2^+^ cells and comparing to other phenotypes, the CRC Metacohort scRNAseq tumor cell data was pseudobulked to avoid false discoveries associated with the large sample size of scRNAseq data. Since there was significant batch effect seen between cohorts, the SCT batch corrected counts were extracted and used for pseudobulking. As such, pseudobulk matrix was generated by aggregating cells for each patient and summing gene expression counts across the sample. Pseudobulked counts were processed with DESeq2 to obtain VST normalized expression values. Combining the signatures compiled by Qin et al. with additional signatures of interest, a list of various stem cell phenotype signatures reported in CRC was generated for analysis (**Supplementary Table 4**). Enrichment scores were computed for each gene signature (n = 75) per patient sample with the GSVA package. The results were used to generate a Pearson correlation matrix; signatures without a significant correlation to gsASCL2 were removed and results visualized with the corrplot package.

### MAPKi Treated PDX RNAseq Analysis

Data from previous CRC PDX model studies were compiled, including data for C5002, C5003, C8120, C0999, C1047, B8131, and C1138. Mutation status for genes of interest (KRAS, BRAF, etc) were obtained for each model from internal PDX database and included as binary (mutated, or WT) for heatmap. Specific MAPKi treatment combination used in the corresponding study was also annotated for each dataset in the heatmap. RNAseq raw counts were obtained for each sample and processed using DESeq2 for each model, with at least three replicates for each group. A list of DEGs between the MAPKi and vehicle group was produced, and genes ranked using the Wald statistic for subsequent GSEA. MAPK signaling was assessed using the published MAPK Pathway Activity Score (MPAS)^86^ gene signature and the resulting normalized expression scores (NES) recorded per model. DESeq2 normalized ASCL2 gene expression values were obtained for each sample; values of MAPKi-treated samples were divided by the average expression of vehicle samples and log transformed (log2FC). Molecular and treatment data for each model were compiled into a summary heatmap along with MPAS NES and ASCL2 log2FC values.

### Generating and Culturing PDX-derived Organoids

With a 0.5-1.0 mm^3^ piece of a PDX tumor, the tissue was minced and dissociated for 2 hours at 37°C using Advanced DMEM/F12 (Corning), 10 mM HEPES (Thermo Fisher), 1x GlutaMAX (Thermo Fisher), 5 mg/mL collagenase II (Thermo Fisher), 1 mg/mL dispase II (Thermo Fisher), and 2.5% FBS (Corning). Dissociated cell mixture was passed through a 40 um cell strainer, pelleted by centrifugation at 300-400xg for 5 minutes at 4°C, and washed three times with wash buffer: Advanced DMEM/F12 (Corning), 10 mM HEPES (Thermo Fisher), 1x GlutaMAX (Thermo Fisher), and 2.5% FBS (Corning). Resulting cell pellet was resuspended with Matrigel (Corning), plated on 24-well plates, and placed in a 37°C incubator after media was added. PDOs were passaged every 2 weeks or when confluent. Organoid growth media (OGM) was changed every 3-5 days and consisted of: Advanced DMEM/F12 (Corning), 10 mM HEPES (Thermo Fisher), 1x GlutaMAX (Thermo Fisher), 1x Wnt3a-conditioned media, 500 nM A83-01 (Thermo Fisher), 50 ng/mL recombinant human EGF (R&D), 100 ng/mL rhNoggin (R&D), 100 ng/mL rhFGF10 (R&D), 10 nM gastrin I (Tocris), 1.25 mM n-acetylcysteine (Sigma Aldrich), 10 mM nicotinamide (Sigma Aldrich), 1x B-27 supplement (Gibco), 10.5 uM Y27632 (Tocris), and 1x Primocin (InvivoGen).

### PDO *Ex Vivo* Assay

C1047 (KRASmut) and B8219 (KRASmut) PDOs were plated using 24-well plates (Corning) and OGM then treatment with appropriate agents based on model. C1047 was treated with vehicle (DMSO) or BI3406 (4uM) plus trametinib (10nM). The MAPKi treatment arm of B8219 received AMG510 (1uM) and cetuximab (50 ug/mL). Plates were imaged with a Cytation 7 (BioTek) every 12 hours for a week, using 10x brightfield z-stack images. Produced images were processed using Gen5 software (BioTek) to generate z-projections, and analyzed to calculate the sum organoid area per well at each timepoint, and values were normalized to the sum organoid area at Day 0. Each well was used a biological replicate, and a minimum of 3 replicates per group were included. Organoid area percent change values were plotted with Prism (GraphPad) and the last timepoints used for a two-tailed t-test for statistical significance. After treatment and monitor period was completed, organoids were collected using Gentle Cell Dissociation Reagent (Stemcell Technologies) and processed for downstream analysis.

### MAPKi Treated PDC Experiments

Reagent concentrations used with each cell lines were determined by cell viability assays to generate dose response curve. Assay was performed using CellTiter Glo (Promega) and a Synergy neo2 (BioTek) plate reader. Absorbance values were normalized to vehicle group and nonlinear regression performed with Prism (GraphPad) to calculate IC50 values. For treatments with two compounds, optimal concentrations for each agent were identified by analyzing relative viability values with SynergyFinder^87^. Media for all cell lines was RPMI (Corning), 10% FBS (Cornign), and 1x Primocin (InvivoGen). Tested PDC models included B8219, C1138 and C1035, each of which were treated with vehicle (DMSO) or MAPKi for 72 hours before samples (n = 3) collected for downstream analysis. MAPKi therapy for B8219 was AMG510 (1uM) and cetuximab (50 ug/mL), and for C1138 and C1035 MAPKi was BI3406 (4uM) and trametinib (10nM).

### Quantitative PCR

Samples of cell pellets were processed using a RNeasy Kit (Qiagen) to extract RNA and yield quantified by Nanodrop One spectrophotometer (Thermo Fisher). The cDNA was generated with iScript Reverse Transcription Supermix (BioRad) and the PowerTrack SYBR Green Master Mix (Thermo Fisher) was used for the qPCR reaction. PrimeTime qPCR Primers (IDT) were purchased for genes of interest, including ASCL2 and ACTB, and sequences for all primers used listed in **Supplementary Table 1**. For the 16 gene qPCR panel, the queried genes included ASCL2, stem-related genes (SOX2, NANOG, SOX9, POU5F1 [OCT4], KLF4, SNAI1), stem-cell marker genes (PROM1 [CD133], CD44, EMP1, LGR5, LRIG1), differentiation markers (CDX2, KRT19, EPCAM), and ACTB as the reference gene. A CFX96 Touch Deep Well Real-Time PCR (BioRad) was used and resulting Cq values were obtained using CFX Maestro software (BioRad).

### Histology Staining and Quantification

Archival FFPE blocks of prior PDX *in vivo* studies conducted in the Kopetz lab were submitted to the DVMS Veterinary Pathology Services core facility (MDACC) for sectioning and staining. Slides were deparaffinized using xylene and ethanol before treatment with EDTA pH 8 retrieval solution (Abcam) at 95°C for 15 minutes. Slides were incubated with Bloxall to quench peroxidase activity and then blocked with 2.5% goat serum (Vector Laboratories) for 1 hour. After washing with TBST buffer, slides were incubated with anti-ASCL2 polyclonal rabbit antibody (PA5-41489, Thermo Fisher) at 1:1000 overnight at 4°C. Anti-rabbit HRP conjugated secondary antibody (Cell Signaling Technology) was added to slides and incubated for 30 minutes before adding SignalStain DAB chromogen (Cell Signaling Technology). Slides were counterstained with hematoxylin, dehydrated with ethanol and xylene, and cover slipped using Cytoseal60 mounting medium (Thermo Fisher). Slides were imaged using an Aperio AT2 system (Leica) at 20x and analyzed using HALO.

### ASCL2^GFP^ and ASCL2^DTR/RFP^ Models

Full list of all lentivirus models and corresponding designs are outlined in **Supplementary Table 2**. To visualize ASCL2^+^ cells we used the stem cell ASCL2 reporter (STAR) system generated by Oost et al.^36^ based on its ability to denote ASCL2 activity in human and murine models, which we refer to as ‘ASCL2 promoter’ (ASCL2p). For the ASCL2^GFP^ system, a lentivirus vector was designed with the ASCL2 promoter driving mNeonGreen (referred to as ‘GFP’ for simplicity), and included a tagBFP and hygromycin resistance cassette for selection {ASCL2p-mNeonGreen, CMVp-tagBFP::Hygro^R^}. The ASCL2^GFP^ lentiviral plasmid and packaged virus were purchased from VectorBuilder (VB220420-1193rct). The ASCL2^DTR/RFP^ system similarly used a lentivirus vector with the ASCL2 promoter driving expression of diphtheria toxin receptor (DTR)^30,59^ and tdTomato (termed ‘RFP’ for simplicity), and a universally expressed puromycin resistance gene to enable antibiotic selection {ASCL2p-DTR:T2A:tdTomato, CMVp-Puro^R^}. The ASCL2^DTR/RFP^ vector and packaged lentivirus were purchased from VectorBuilder (VB221213-1289nzd). Additionally, an ASCL2^DTR/GFP^ system was produced by switching the tdTomato (RFP) gene for mNeonGreen (‘GFP’), and changing the selection method to a tagBFP and hygromycin resistance cassette {ASCL2p-DTR:T2A:mNeonGreen; CMVp-tagBFP:T2A:Hygro^R^}. The ASCL2^DTR/GFP^ vector and lentivirus were obtained from VectorBuilder (VB231207-1264etu). All cell lines were transduced with the indicated lentivirus at an MOI of 5, with 8 ug/mL Polybrene (VectorBuilder) added to standard cell media. Response curves with parental cell lines were generated for puromycin (Gibco) and hygromycin (Gibco), as described above, to determine their IC_90_ concentrations. Transduced cell lines were treated with the appropriate selection antibiotic for three passages. ASCL2^GFP^ and ASCL2^DTR/GFP^ cells were flow-sorted using a FACSAria Fusion (BD) cytometer (MDACC Flow Cytometry and Cellular Imaging Core Facility) to collect the BFP^+^ cell population for further use. Models were generated using the following cell lines: B1003 (BRAF^V600E^), WiDr (BRAF^V600E^), DLD1 (KRAS^G12D^), CT26 (BRAF/KRAS^WT^), and MC38 (BRAF/KRAS^WT^). Additionally, the ABPS murine CRC cell line, which has APC, TP53, SMAD4, and BRAF^V600E^ mutations, was obtained from the Corcoran Lab (Massachusetts General Hospital)^16^ and also used to generate models. ABPS cells were cultures with DMEM/F12 media (Corning), 10% FBS (Corning), and 1x Primocin (InvivoGen). As negative controls, cell lines were similarly generated with a control lentivirus which expressed mCherry and EGFP {EF1Ap-mCherry, CMVp-EGFP:T2A:Puro^R^} (VB010000-9298rtf).

### ASCL2^+^ Flow Cytometry

B1003 and DLD1 cell lines transduced with ASCL2^GFP^ lentivirus, as outlined above, were flow-sorted using a FACSAria Fusion (BD) by GFP expression. Collected GFP^+^ and GFP^-^, or GFP^High^ and GFP^Low^, cell populations (2x10^6^ cells) for each model were pelleted and cells used for subsequent experiments. Ablation efficacy of ASCL2^+^ cells using DTR system was tested using CT26 ASCL2^DTR/RFP^ model. After 24 hours from being plated, cells were treated with vehicle (PBS) or 100 ng/mL DT (Sigma Aldrich); triplicate samples of each treatment were collected every 24 hours for 3 days. At each collection timepoint, cells were trypsinized, washed with PBS, and fixed overnight with 2% paraformaldehyde (PFA) at 4°C; cells were then washed with PBS and stored in freezing media. For analysis, samples were washed, resuspended in flow buffer (PBS with 1% BSA), and processed using a LSRFortessa X-20 Cell Analyzer (BD). Results were analyzed using FlowJo, where ASCL2^+^ cell prevalence was calculated as the number of RFP^+^ cells over the total number of gated cells in the sample, and a two-way ANOVA test was performed with post-hoc Bonferroni for multiple comparison. Similarly, to test ASCL2^+^ ablation efficacy in MAPKi-treated cells, CT26 ASCL2^DTR/RFP^ samples were processed as before but treatment groups were vehicle (DMSO)/vehicle (PBS), vehicle/DT, MAPKi/vehicle, and MAPKi/DT. For MAPKi treatment, cells were treated with 2 uM BI-3406 and 40 nM trametinib, and DT was dosed at 100 ng/mL.

Remainder of methods were as described above. Lastly, CRC models transduced with ASCL2^GFP^ lentivirus, including B1003, DLD1, WiDr, and CT26, were used to determine impact of therapy on ASCL2^+^ (APT) cells. Samples for each model were plated and treated with vehicle (DMSO) or a MAPKi regimen depending on the mutation status of the individual model. B1003 and WiDr received encorafenib (BRAFi) and cetuximab (EGFRi) for MAPKi, while the other models were treated with BI-3406 (SOSi) and trametinib (MEKi). All therapeutic agents were used at previously determined IC_50_ concentrations based on cell viability assays, as described previously. Samples for each model were collected after 24, 48 and 72 hours of treatment and processed for flow cytometry analysis. After being collected, cells were fixed with 2% PFA overnight at 4°C, then resuspended in flow buffer before processing with a LSRFortessa X-20 Cell Analyzer (BD). As before, data was analyzed with FlowJo to determine ASCL2^+^ (GFP^+^) cell prevalence in each sample, and results compared using a two-way ANOVA with post-hoc Bonferroni for multiple comparison correction.

### Immunofluorescent Live-Cell Imaging

Cells from B1003 ASCL2^GFP^ model were plated after flow-sorting by GFP expression, as outlined above, with a FACSAria Fusion (BD). GFP^High^ and GFP^Low^ cells were plated in 96-well black plates (Corning) and incubated at 37°C in an Incucyte S3 (Sartorius). Brightfield phase contrast and GFP fluorescent images of cells were collected every 12 hours for 5 days, and images processed with standard Incucyte cell analysis module. To determine ablation efficacy of DTR system, CT26 ASCL2^DTR/RFP^ cells were stained with CellEvent Caspase 3/7 Green Detection Reagent (Invitrogen) to visualize apoptotic cells. Samples using parental CT26 cells served as negative control while WiDr cells were used as a positive control. After treatment with vehicle (PBS) or 100 ng/mL DT, cells were imaged using an Incucyte (Sartorius) with phase contrast, GFP and RFP images collected every 4 hours for 3 days. Images were processed with standard analysis module, and the ASCL2^+^ cell ablation efficacy was computed using the fraction of apoptotic ASCL2^+^ cells (GFP^+^/RFP^+^ cell area) relative to the total ASCL2^+^ cell population (RFP^+^ cell area) for each sample (n = 6). Results were graphed and 72 hour data used to perform a t-test for statistical significance. The same procedure was used for DT ablation with MAPKi treatment experiment with the following exceptions. Treatment groups included vehicle (DMSO)/vehicle (PBS), vehicle/DT, MAPKi/vehicle, and MAPKi/DT, with MAPKi treatment consisting of 2 uM BI-3406 and 40 nM trametinib; 100 ng/mL DT was used. Samples were processed similarly but AUC values calculated per treatment group, and a one-way ANOVA with post-hoc Tukey test was used for significance. For cell growth assays, after cells were plated, following same procedure, phase contrast images were collected every 12 hours for 5-7 days using an Incucyte (Sartorius). Resulting images were processed using Incucyte analysis module to identify cells in each well and the resulting confluence per well values were used to measure cell growth over time as confluence (%) or normalized confluence. Data was graphed and AUC values utilized for a one-way ANOVA with post-hoc Tukey for multiple comparison (n = 6). This methodology was used for both CT26 ASCL2^DTR/RFP^ and ABPS ASCL2^DTR/GFP^, where MAPKi treatment of CT26 was as listed above, and for ABPS MAPKi consisted of 1 uM encorafenib (BRAFi), 300 uM cetuximab, and 10 nM trametinib.

### ASCL2^DTR/RFP^ Cell Viability Experiments

CT26 ASCL2^DTR/RFP^ and ABPS ASCL2^DTR/GFP^ models were used to see impact of DT ablation of ASCL2^+^ (APT) cells on MAPKi therapy efficacy. Parental CT26 and ABPS cell lines were also tested to evaluate untransduced models. As such, each model was plated in 96-well black plates (Corning) and treatment added 24 hours after cells were plated. Samples were treated with vehicle (PBS), 100 ng/mL DT, MAPKi, or MAPKi/DT, incubated at 37°C, and one plate collected and assayed at 24, 48 and 72 hours. MAPKi treatment for CT26 samples was 2 uM BI-3406 and 40 nM trametinib, whereas ABPS received 1 uM encorafenib, 300 uM cetuximab and 10 nM trametinib. At each collection, CellTiter Glo (Promega) was used with a Synergy neo2 (BioTek) plate reader to evaluate relative viability of each sample relative to vehicle. Data was graphed using Prism (GraphPad) and a two-way ANOVA with post-hoc Bonferroni (n = 6) used for statistical analysis.

### APT Cell MAPKi Response Assay

Sensitivity of APT (ASCL2^+^) cells to MAPKi therapy relative to other CRC cells (ASCL2^-^) was evaluated by cell viability assay using flow-sorted the CT26 ASCL2^DTR/RFP^ model. Previous results for CT26 showed IC_50_ values of 2 uM BI-3406 and 40 nM trametinib, therefore we treated all cells with 1 uM BI-3406 and varied the trametinib concentration (0-1 uM). Cells were flow-sorted using a FACSAria Fusion (BD) by RFP expression to collect RFP^High^ (‘High’) and RFP^Low^ (‘Low’) cell populations, as detailed above. Three cell groups were tested using the collected cells: RFP^High^ cells without DT, denoted as High/-DT, Low/-DT, and Low/+DT. For DT treatment, samples received either vehicle (PBS) or 100 ng/mL DT. After 72 hours of incubation at 37°C, plates were processed with CellTiterGlo (Promega) and absorbance read with a Synergy neo2 (BioTek) plate reader to determine the relative viability. Results were graphed with Prism (GraphPad), nonlinear regression performed to determine the trametinib IC_50_ values for each group, and compared using a one-way ANOVA with post-hoc Bonferroni.

### Spheroid Forming Limited Dilution Assay

CT26 ASCL2^DTR/RFP^ cells were sorted using a FACSAria Fusion (BD) to collect RFP^High^ and RFP^Low^ cell populations. An *in vitro* limited dilution assay was conducted, as previously described by Agro et al.^88^, to determine the stem capacity of ASCL2^+^ (APT) and ASCL2^-^ cells. RFP^High^ and RFP^Low^, referred to as ‘High’ and ‘Low’, were ran in duplicate with one receiving vehicle (PBS) and the other 100 ng/mL DT. Spheroid media consisted of DMEM/F12 (Corning), 10 mM HEPES (Thermo Fisher), 1x GlutaMAX (Thermo Fisher), 50 ng/mL human EGF (R&D), 100 ng/mL human FGF10 (R&D), 1x B-27 supplement (Gibco), and 1x Primocin (InvivoGen). The flow-sorted cells were serially diluted to obtain 6 suspensions of varying cell densities to achieve the desired number of cells per well when plated. Cell suspensions in spheroid media were plated using Nuclon 96-well U-shaped ultra-low attachment plates (Thermo Scientific) as follows: 1000 cells/well (n = 24 wells), 500 cells/well (n = 24), 100 cells/well (n = 48), 50 cells/well (n = 48), 10 cells/well (n = 96), and 1 cell/well (n = 192). Plates were incubated at 37°C and monitored for spheroid formation over the course of two weeks. Number of wells that successfully formed a spheroid was collected for each group, and data was processed using the Extreme Limiting Dilution Analysis (ELDA) software (http://bioinf.wehi.edu.au/software/elda/) to determine stem-cell frequency and perform statistical analysis. Similar procedure was used for WiDr ASCL2^GFP^ cells which were flow-sorted by GFP expression and used to determine stem-cell frequency of GFP^High^ and GFP^Low^ cell populations.

### Colony Formation Assay

CT26 ASCL2^DTR/RFP^ cells were used to determine the colony formation efficacy of a standard cell population, representing mixture of APT cells and normal CRC cells, compared to one depleted of APT (ASCL2^+^) cells. CT26 ASCL2^DTR/RFP^ cells were plated in a 10 cm petri dish with RMPI media with 10% FBS and treated with vehicle (PBS) or 100 ng/mL DT for 48 hours, before being collected and used to seed 500 cells/well in 12-well plates for each group. Plates were monitored for 1-2 weeks for growth, with media changed every 5 days, and were imaged using an Incuctye (Sartorius) to collect whole well brightfield images. The number of colonies in each well were counted manually and the colony formation rate was computed by dividing the number of counted colonies by the 500 cells that were seeded. Colony formation efficiency was calculated by normalizing the formation rate to the rate of the vehicle group. Results were graphed and a two-sided t-test was performed.

### ASCL2 Knockdown and Overexpression

ASCL2 knockdown (KD) was performed in DLD1 cells using a TriFECTa DsiRNA kit (IDT) with three ASCL2 DsiRNAs, referred to as A1-A3, and a non-coding (NC) control (**Supplementary Table 1**). Cells were transfected using the 0-10 nM DsiRNAs and Lipofectamine RNAiMAX (Thermo Fisher Scientific), as per the manufacturer instructions using Opti-MEM reduced serum medium (Gibco). Samples were collected after 72 hours of treatment and qPCR performed, as outlined above, to determine ASCL2 KD efficiency, which found 10nM A2 DsiRNA produced highest ASCL2 silencing and was used for subsequent studies. Impact of ASCL2 KD on MAPKi treatment was tested by monitoring cell growth of DLD1 cells treated with 10nM A2 or NC DsiRNA, and received vehicle (DMSO) or MAPKi (SOSi and MEKi) therapy. Treated cells were imaged every 6 hours for 3 days using an Incucyte (Sartorius), and images processed using standard analysis module to determine confluency of wells at each timepoint. Results for each treatment group (n = 3) were plotted and data from last timepoint used to perform a one-way ANOVA with Bonferroni multiple comparison correction. To evaluate impact of overexpressing ASCL2 in CRC cells, a lentiviral vector was designed using the CMV promoter to drive expression of human ASCL2 gene (NM_005170.3) and with a neomycin resistance cassette to enable antibiotic selection {CMVp-hASCL2; mPGKp-NeoR} (VB230901-1318gpz). The ASCL2 overexpression packaged lentivirus was obtained (Vectorbuilder) and used to transduce WiDr and B1003 cells. After three rounds of selection with neomycin, models were used for subsequent analysis and labelled as ‘hAO’ or ASCL2^OE^ models. ASCL2 overexpression was confirmed by qPCR and western blot. WiDr hAO cells were used to conduct cell growth assay and colony formation assay, as previously described. Briefly, parental (WT) and hAO WiDr cells were plated, treated with vehicle or MAPKi (encorafenib, cetuximab), and imaged every 12 hours for 3 days using an Incucyte (Sartorius) to determine well confluency relative to starting confluency at day 0. For the clonogenic assay, parental (WT) and hAO cells were plated and number of colonies per well counted after 7-10 days, with results reported as colony formation efficiency relative to parental model.

### Generating Molecular Barcoded B1003b Model

B1003 (BRAF^V600E^) was used to generate a barcoded PDX model of CRC as follows. The B1003 PDX-derived cell line was cultured using RPMI media with 10% FBS and 1x Primocin (InvivoGen). The CloneTracker XP 10M Barcode-3’ library with Venus-Puro (Cellecta) was used according to user manual and methods described by Seth et al.^89^ in order to barcode the B1003 cells using an MOI of 0.3. Transduced cells underwent antibiotic selection using Primocin (Gibco) and were then passaged multiple times to expand and collect samples for analysis. To determine the model was properly barcoded and obtain barcode IDs of the established cells, the NGS Prep Kit for Barcode Libraries in pScribe (Cellecta) was used according to the manufacturer instructions. Resulting samples were then submitted for sequencing (HiSeq) and FASTQ files processed using the NGS Demultiplexing and Alignment Software (Cellecta). This approach of identifying barcode IDs for a sample is referred to as BarcodeSeq throughout this work. The established B1003 barcoded model is denoted ‘B1003b’, and additional experiments using the model are described below.

### Mouse *In Vivo* Studies

All murine studies were conducted in accordance with ethical regulations and institutional policies as approved by the University of Texas MD Anderson Cancer Center Institutional Animal Care and Use Committee (IACUC). Mice received water and standard diet *ad libitum*, unless otherwise specified, and were housed in a light-dark cycle, temperature controlled facility. Methods used for previously established PDX models are described elsewhere^90^. Murine studies used 6-8 week old female athymic nude mice (Envigo), which were sedated with isoflurane and subcutaneously injected with 200uL of CRC cell suspension in the flank. Each of the used cell models was resuspended in PBS and diluted at 1:1 ratio with Matrigel (Corning) before being injected. Mice were monitored daily to assess health and tumor volume recorded every 2-3 days. Tumors were measured by recording the longest dimension as ‘length’ (L) and the second dimension as ‘width’ (W), to estimate tumor volume using (L x W^2^)/2 in mm^3^ units. Mice were randomized into treatment groups when tumors reached approximately 250 mm^3^ in volume and were monitored daily for the duration of the study. Additional details will be described in their respective section for each mouse study. For the barcoded B1003 (B1003b) model, 5x10^6^ cells were injected to generate subcutaneous tumors. Treatment groups in this study were the vehicle and MAPKi groups, each of which had 5 mice. Dosing of treatments for this study are outlines in ‘Pharmacological Treatments’ below. After 21 days of treatment mice were euthanized and B1003b tumors were collected for subsequent analysis including scRNAseq.

### Pharmacological Treatments

Mouse study for B1003b used encorafenib and cetuximab as MAPKi therapy. Encorafenib (Selleckchem) was given at 10 mg/kg, twice daily by oral gavage (OG), and cetuximab (MDACC) administered at 20 mg/kg twice weekly by intraperitoneal injection (IP). In CT26 studies, MAPKi treatment consisted of BI-3406 and trametinib. BI-3406 (MedChem Express) was given at 50 mg/kg, twice daily by OG, and trametinib (MedChem Express) administered at 0.25 mg/kg, twice daily by OG. For ABPS studies, a combination of encorafenib, trametinib, and cetuximab were used for the MAPKi treatment groups. Dosing of encorafenib and trametinib was done using specialized mouse diet, consisting of irradiated Purina Rodent Diet (#5053) with 110 ppm encorafenib and 2.8 ppm trametinib (Research Diets, Inc.). The MAPKi chow was given to treatment group mice *ad libitum*, and cetuximab (MDACC) was administered at 20 mg/kg twice weekly by IP injection. For all studies containing mice treated with DT, additional measures outlined by biohazard safety risk protocol were used as precaution, including housing mice in biohazard unit. DT (Sigma Aldrich) was dosed at 50 ug/kg and administered by IP injection every other day.

### scBarcodeSeq Workflow and Data

B1003b tumors from mouse study were collected as described above. Tumors were dissociated into a single cell suspension using the methods outlined in the generating organoids section above. Briefly, after mincing the tissue a mixture of collagenase/dispase was used to dissociate the tumor, and the final cell pellet was resuspended in sequencing buffer (PBS + 1% BSA). Samples were processed by the Single Cell Core (MDACC) using 10x Genomics 3’ Chromium V3 and sequenced using Novaseq 6000 (Illumina). FASTQ files were processed with CellRanger (10x Genomics) for read alignment to the human (GRCh38) and mouse (GRCm38) genomes. CellRanger outputs from each sample were used to generate individual Seurat objects and then merged together using the Seurat v5 package.

Identification of barcode IDs in the sequenced cells was performed using the following workflow, which is based on manufacturer (Cellecta) recommendations in CloneTracker XP barcode library manual. Molecular barcodes are DNA integrated and mRNA expressed in the polyA tail of the venus gene, which has low expression in cells and is difficult to obtain through standard scRNAseq workflow. Custom primers can be used with the cDNA generated from 10x Chromium 3’ processing to amplify barcodes (referred to as ‘barcodeID’) along with the corresponding 10x cell barcode (termed ‘cell ID’), enabling pairing of barcode IDs with matching cells in scRNAseq data based on cell IDs. The custom primers were designed (**Supplementary Table 1**) and purchased from IDT. As such, half of the cDNA produced from scRNAseq of each sample was used as follows. Sample cDNA was combined with 300 nM P5_Read1 primer, 300 nM P7_adapter primer, 1x dNTPs (Cellecta), 1x taq buffer (Cellecta), and 1x taq polymerase (Cellecta). Samples were run on a CFX96 Touch Deep Well Real-Time PCR (BioRad) using the following conditions: 1) 95°C for 2 minutes 2) 95°C for 30 seconds, 3) 65°C for 30 seconds, 4) 68°C for 2 minutes, 5) repeat to #2 for 17 cycles, 6) 68°C for 2 minutes, and 7) hold at 4°C. After the PCR, samples were run on a 1% agarose gel to confirm the produced DNA was the expected length (497 bp) and purify the product by gel extraction using a QIAquick Gel Extraction Kit (Qiagen). Yield was assessed with a NanoDrop One spectrophotometer (Thermo Fisher) and samples submitted to the Advanced Technology Genomics Core (MDACC) for 150 PE sequencing with iSeq (Illumina).Resulting FASTQ files were processed using a custom pipeline designed to extract the paired barcode data.

Read1 and read2 FASTQ files were read into R using the CellBarcode package, and reads with poor quality were identified and filtered out using the ShortRead package. The CellBarcode package was used to extract the information from remaining reads including readID and cell ID sequence from read1 using the pattern: ‘([ACTG]{16})([ACTG]{12})TTTTTTTTTTTTTTTTTTTTTTTTTTTTT’. The same process was repeated for read2 using the pattern ‘CCGACCACCGAACGCAACGCACGCA([ATCG]{14})TGGT([ATCG]{30})’ to obtain the 48 bp barcodeID sequence, and matched to corresponding cell ID by readID. BarcodeID sequences were renamed using barcodeIDs (e.g. F035T00768), which are easier to use in downstream analysis. To incorporate the barcodes in the scRNAseq data, the final list of cell IDs and matching barcodeIDs were then added to the metadata layer of the Seurat objects, and further processed for scBarcodeSeq analysis.

### B1003b scBarcodeSeq Analysis

B1003b scRNAseq data combined with the barcode data generated above, is referred to as the scBarcodeSeq dataset. QC filtering of cells was performed to remove cells with percent mitochondria gene count > 10% and gene count (nFeature_RNA) < 200. The SCTransform normalization workflow was used to normalize counts, identify variable genes, and scale gene values, and a UMAP was generated using FindUMAP function to visualize the data. To identify the tumor cell population, the FeaturePlot function was utilized to visualize the percent mouse reads per cell to identify cell clusters corresponding to mouse (stroma, immune, etc.) and human (tumor) cells. Given that barcoding was conducted in vitro, only human cells should contain barcodes, therefore data was also visualized by barcodeID to see if the mouse clusters overlapped with non-barcoded cells. Clusters with high mouse read percent and without barcodes were filtered out. Remaining barcoded tumor cells were reprocessed using SCTransform function to normalize and scale counts, and data was integrated using FastMNN to account for treatment impact and produce a corrected UMAP for visualization. Gene signature values were computed for MPAS, gsASCL2, and other signatures of interest using the AddModuleScore function, and z-scores were calculated for each. Cells with matching barcodeIDs in vehicle and MAPKi samples were identified and labeled as ‘replicate barcodes’. Since some replicate barcodes had multiple cells with the same barcodeID within a sample, MPAS and gsASCL2 scores were averaged by barcodeID and treatment, before conducting the paired analysis between vehicle and MAPKi using a paired t-test. Cells in the dataset were classified as ‘ASCL2 positive’ and ‘ASCL2 negative’ using the z-scored gsASCL2 values. ASCL2^+^ cell prevalence was computed by dividing the number of ASCL2^+^ cells by the total number of cells in the sample.

### EGFRi Rechallenge Clinical RNAseq Data

Raw gene counts for patient samples (n = 51) were provided by Christine Parseghian (MDACC) from clinical trials NCT03087071 and NCT04616183; clinical annotations included response based on RECIST, patient age, and time-to-progression (TTP) in days. In this trial, BRAF/KRAS^WT^ patients with metastatic CRC and prior history of EGFRi therapy were rechallenged with EGFRi therapy to see if they obtained any benefit. Patients (n = 26) underwent biopsy prior to starting treatment (‘Pre’) and one month post-treatment initiation (‘Post’), which were submitted for bulk RNAseq. After an initial filtering to remove unpaired samples or samples with low quality, 40 samples remained corresponding to 7 nonresponders (NR) and 13 responders (R). The data was analyzed using DESeq2 package and vst normalized gene expression values were produced for each patient sample. Normalized values were used in GSVA, along with the MPAS and gsASCL2 gene signatures, to determine the expression of the gene sets in each sample and were z-transformed. Signature scores were compared between the Pre and Post timepoint, while matching samples by patient ID, using a Wilcoxon matched pair signed rank test. Delta ASCL2 signature expression was calculated by subtracting the ‘Post’ ASCL2 Index score (i.e. gsASCL2) from the ‘Pre’ value for each patient. A Pearson correlation was performed to compare delta ASCL2 signature with TTP.

### MAPKi ASCL2^DTR^ In Vivo Studies

Studies described here largely followed methods outlined in ‘Mouse In Vivo Studies’, with deviations from that approach listed below. For the CT26 ASCL2^DTR/RFP^ study, 1x10^6^ cells were injected subcutaneously to generate tumors. Once tumors reached appropriate size, mice (n = 8) were randomized into the following treatment groups: vehicle, DT, MAPKi, or MAPKi/DT. Dosing of DT and MAPKi was as listed above. This study was conducted to assess survival benefit of APT (ASCL2^+^) cell depletion with MAPKi therapy. As such, mice were monitored daily and were euthanized if they were moribund, developed ulcers, tumor reached maximum size, or if flagged by DVMS due to health concerns. Data was visualized using Prism (GraphPad) and Kaplan-Meier survival analysis was performed. A similar study was repeated using the CT26 ASCL2^DTR/GFP^ and ABPS ASCL2^DTR/GFP^ models following the same process, but with the primary intention of assessing tumor volume differences between treatment groups. Therefore, tumor volume was measured and recorded, as outlined above, and results analyzed using one-way ANOVA with post-hoc Bonferroni multiple comparison correction.

## List of Tables

**Supplementary Table 1: List of qPCR primer sequences Supplementary Table 2: List and description of all lentiviral vectors**

**Supplementary Table 3: Signatures used in BRAFi patient cohort analysis and results Supplementary Table 4: List of stem-like CRC signatures used for characterization**

## Author Contributions

OEV and SK designed study and concepts for project. OEV conducted experiments, compiled results, generated figures, and wrote manuscript. OEV, YX, HT, AM, DP, AA, RM, MP, AHM, HML, CWW, NF, PK, AS, JA, OC, KL, CB, AV, JPS, CP, JRM, RC, SK reviewed manuscript.

---

**Supplementary Figure 1.**
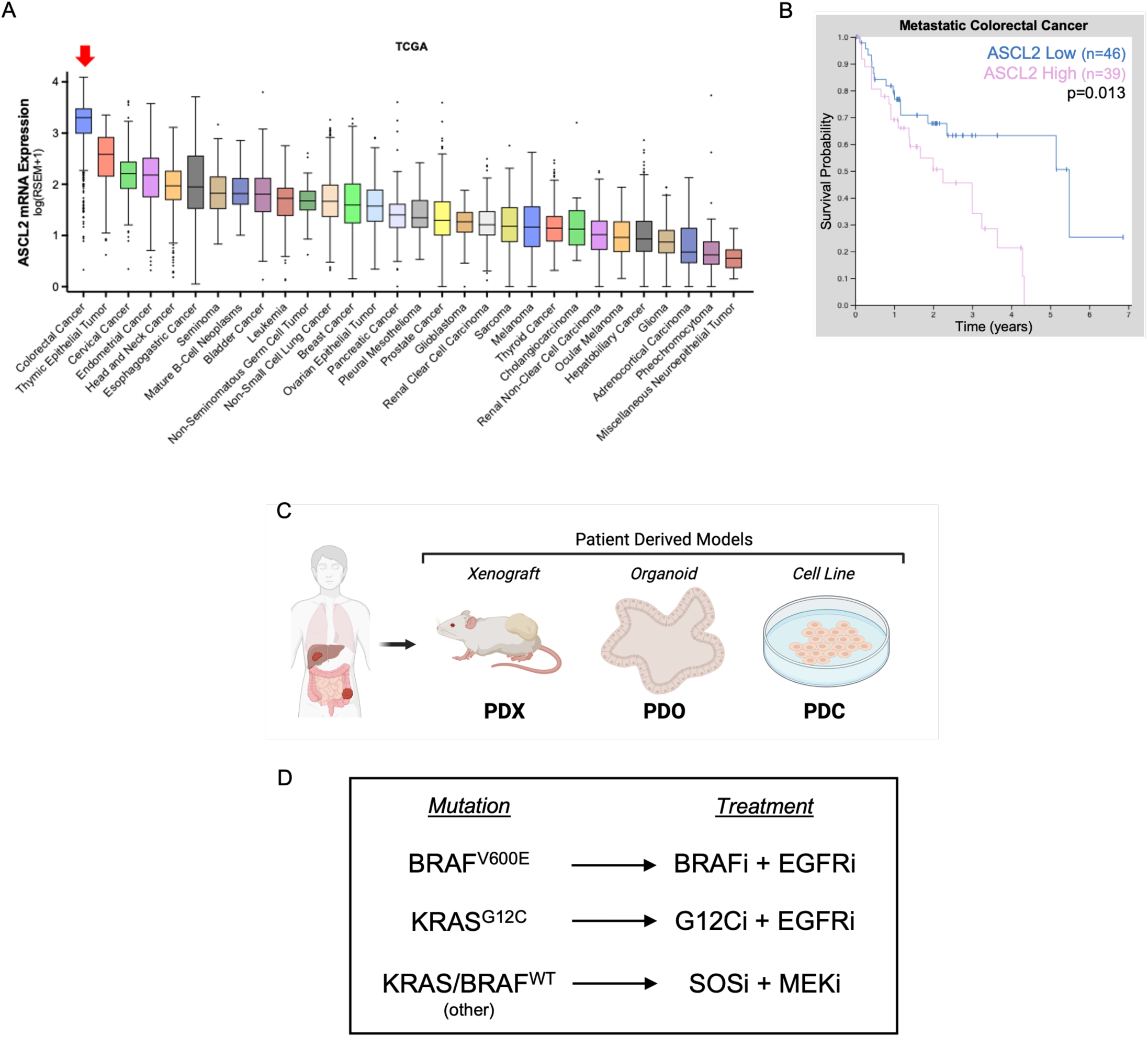
ASCL2 expression in cancer. **A)** Pan-cancer RNA expression of ASCL2 in TCGA dataset. **B)** Kaplan-Meier plot from Human Protein Atlas of patients with mCRC classified based on tumor ASCL2 mRNA expression as either High (n = 39) or Low (n = 46), with log-rank p = 0.013. **C)** Biopsies from patients with CRC are used to generate patient-derived xenograft (PDX), organoid (PDO), and cell lines (PDC). **D)** Overview of MAPKi treatments used based on the individual model’s mutation status, further detailed in methods. Encorafenib was used for BRAFi, cetuximab for EGFRi, sotorasib for G12Ci, BI-3406 for SOSi, and trametinib for MEKi.

**Supplementary Figure 2.**
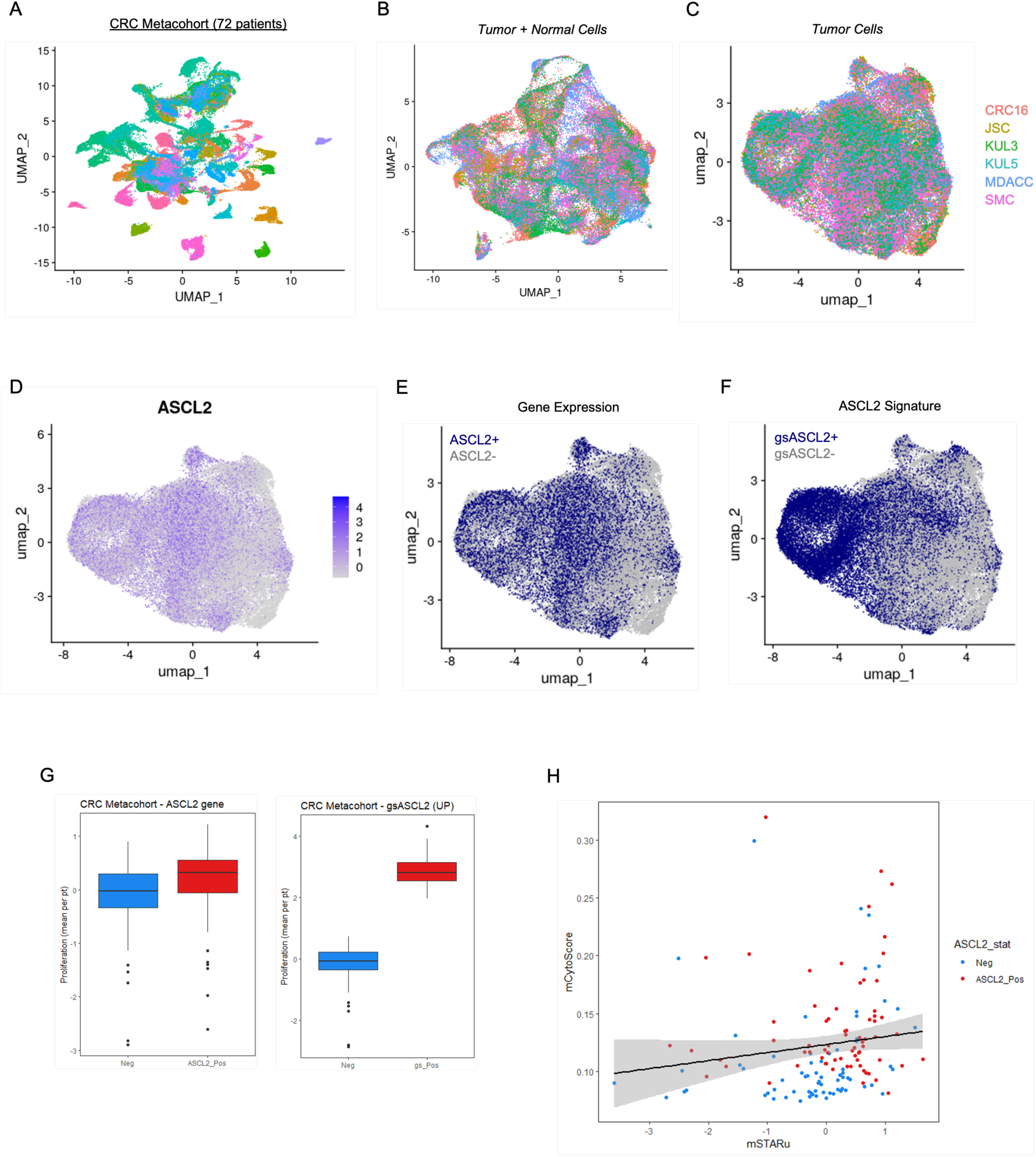
ASCL2 gene signature in CRC metacohort. **A)** UMAP of CRC Metacohort (n = 72) colored by cluster. **B-C)** Batch corrected UMAPs of all cells (**B**) and of only tumor cells (**C**) colored by cohort of the original data. **D)** ASCL2 normalized gene expression in tumor cells of patients. **E-F)** Cells colored by cell phenotype as positive (blue) or negative (grey) based on ASCL2 gene expression (**E**) or gsASCL2 signature expression (**F**). **G)** Average proliferation signature expression per patient from tumor cells classified as negative (blue) or positive (red) by ASCL2 gene (left) or gsASCL2 signature (right) expression. Box plots depict median, first and third quartiles, whiskers reflect 1.5*IQR, and outliers shown as points. **H)** Mean tumor cell expression of gsASCL2 and stem potency per patient, with simple linear regression trendline and 95% CI depicted.

**Supplementary Figure 3.**
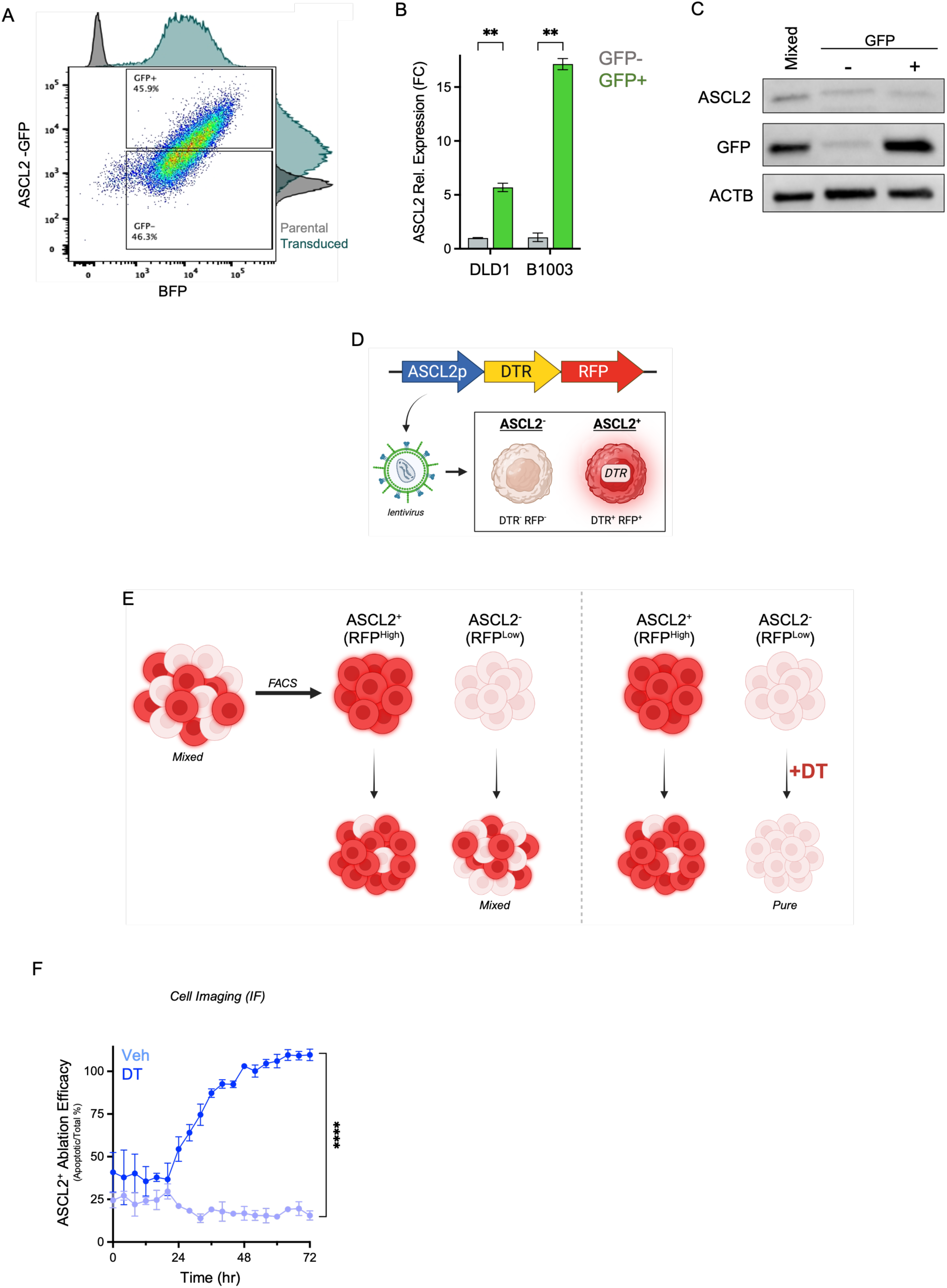
Validation of ASCL2^GFP^ and ASCL2^DTR/RFP^ models. **A-C)** ASCL2^GFP^ models flow-sorted by GFP and BFP expression, collecting GFP^+^/BFP^+^ (ASCL2^+^) and GFP^-^/BFP^+^ (ASCL2^-^) cells. **A)** B1003 flow plot depicting gates defined by single channel histograms of transduced (teal) and parental (grey) cells. **B)** ASCL2 mRNA expression in flow-sorted GFP^+^ (green) cells relative to GFP^-^ (grey) cells for B1003 (p = 0.0013) and DLD1 (p = 0.0021) models (n = 3; two-sided Welch’s t-test; mean ± SEM). **C)** Western blots of B1003 unsorted (mixed) and sorted GFP^+^ and GFP^-^ samples for ASCL2, GFP (mNeonGreen), and ACTB. **D)** Lentiviral ASCL2^DTR/RFP^ or ASCL2^DTR/GFP^ system used for DT induced ablation of ASCL2^+^ cells, which express RFP (or GFP) and DTR (Supplementary Table 2). **E)** Phenotypic plasticity complicates use of cell populations derived from flow-sorted CRC ASCL2^DTR/RFP^ models, as RFP^Low^ samples quickly become phenotypically mixed due to cells acquiring an ASCL2^+^ state. However, DT treatment of RFP^Low^ samples ablates cells that adopt an ASCL2^+^ state, resulting in a homogenous ASCL2^-^ cell population. **F)** APT (ASCL2^+^) cell ablation efficacy of CT26 ASCL2^DTR/RFP^ cells treated with vehicle or DT; cells stained with apoptosis (GFP) marker and samples (n = 6) monitored by IF imaging (AUC and two-sided t-test; mean ± SEM).

**Supplementary Figure 4.**
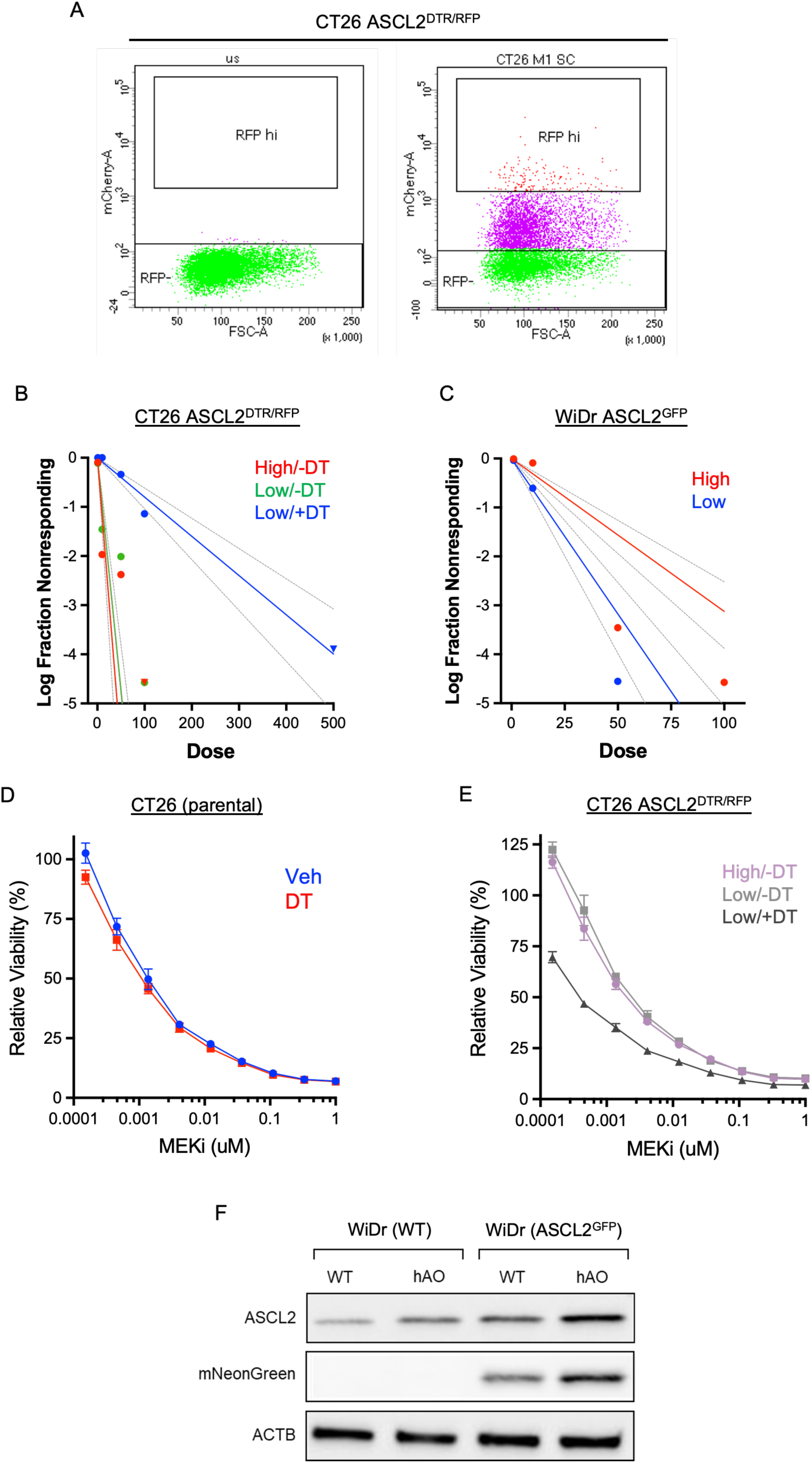
Additional characterization of ASCL2^+^ cells. **A)** Flow-sorting of CT26 ASCL2^DTR/RFP^ model by RFP expression; collection gate defined using parental CT26 (left) cells, and samples of RFP^High^ (termed ‘High’) and RFP^Low^ (termed ‘Low’) were collected (right) for additional study. **B)** Spheroid forming limited dilution assay with sorted CT26 ASCL2^DTR/RFP^ cells, with all three tested groups: High/-DT (red), Low/-DT (green), and Low/+DT (blue); showing no difference in stem-cell frequency between High/-DT and Low/-DT (p = 0.11). **C)** WiDr ASCL2^GFP^ flow-sorted by GFP expression and spheroid forming limited dilution assay conducted using GFP^High^ (High) and GFP^Low^ (Low) cells. **D-E)** Cell viability of CT26 treated with 1uM SOSi (BI-3406) and 0-1uM MEKi (trametinib) to determine IC_50_ values. Control assay performed with parental cells (**D**) treated with vehicle (blue) or DT (red). Flow-sorted CT26 ASCL2^DTR/RFP^ cells (**E**) used to compare response between High/-DT (purple), Low/-DT (light grey), and Low/+DT (grey). **F)** Western blots of ASCL2, GFP (mNeonGreen), and ACTB using WiDr parental (WT) and ASCL2^GFP^ cell lines with ASCL2 overexpression (hAO).

**Supplementary Figure 5.**
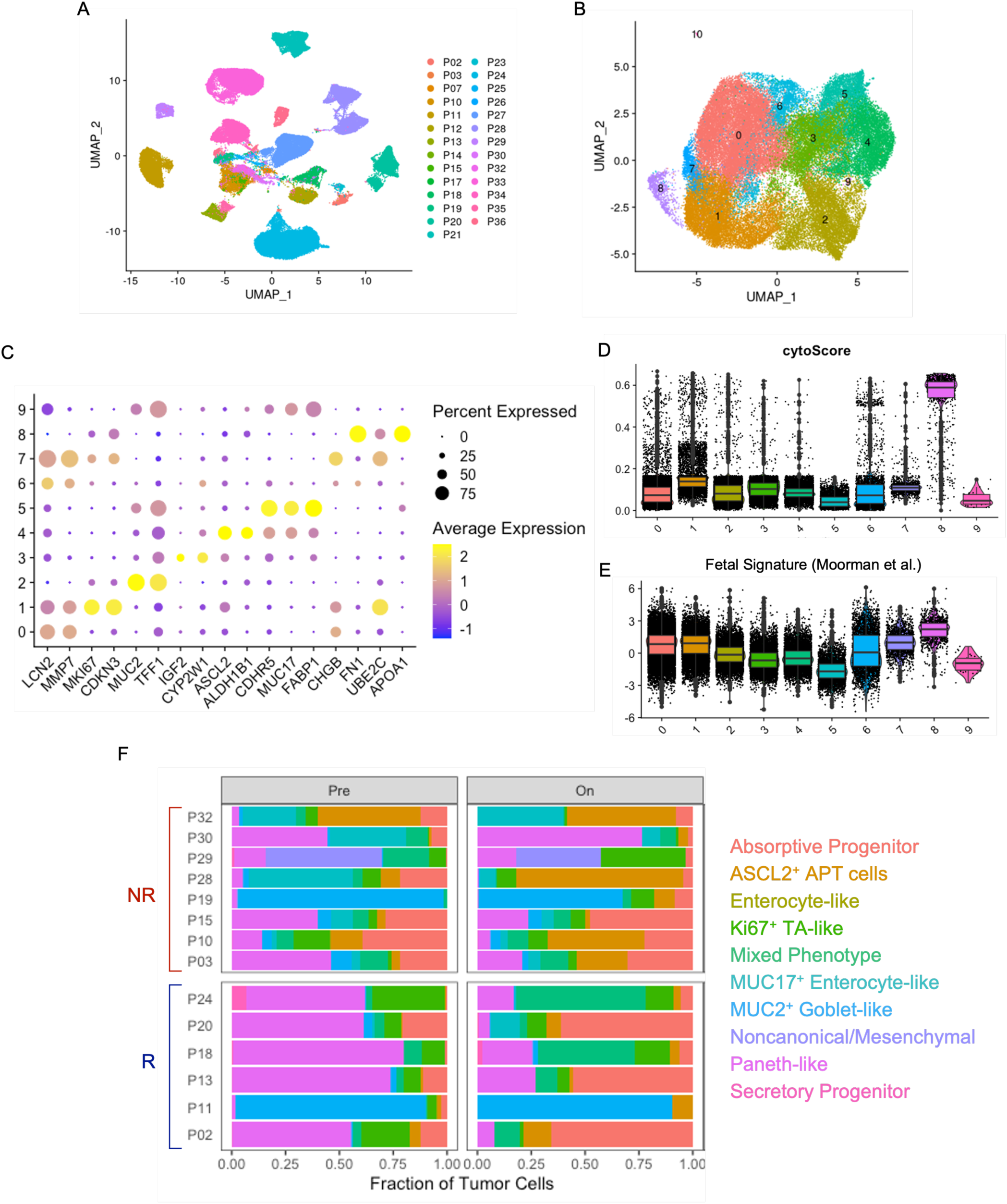
ASCL2 defines a specific cell population in patient tumors. **A)** BRAFi patient cohort scRNAseq data displayed as UMAP with all available samples colored by patient. **B)** Seurat clusters denoting 11 cell phenotypes using batch corrected UMAP of patient (n = 14) sample colored cluster. **C)** Dot plot of genes used to identify cell phenotypes of each cluster; cluster 10 was excluded since they were apoptotic cells. **D-E)** Violin plots with overlapping box plots of the stem potency (cytoScore) and fetal intestinal signature (Moorman et al.) expression by cell cluster used to identify phenotypes. **F)** Stacked bar plot of each patient sample’s tumor cell composition defined by the identified cell cluster phenotypes.

**Supplementary Figure 6.**
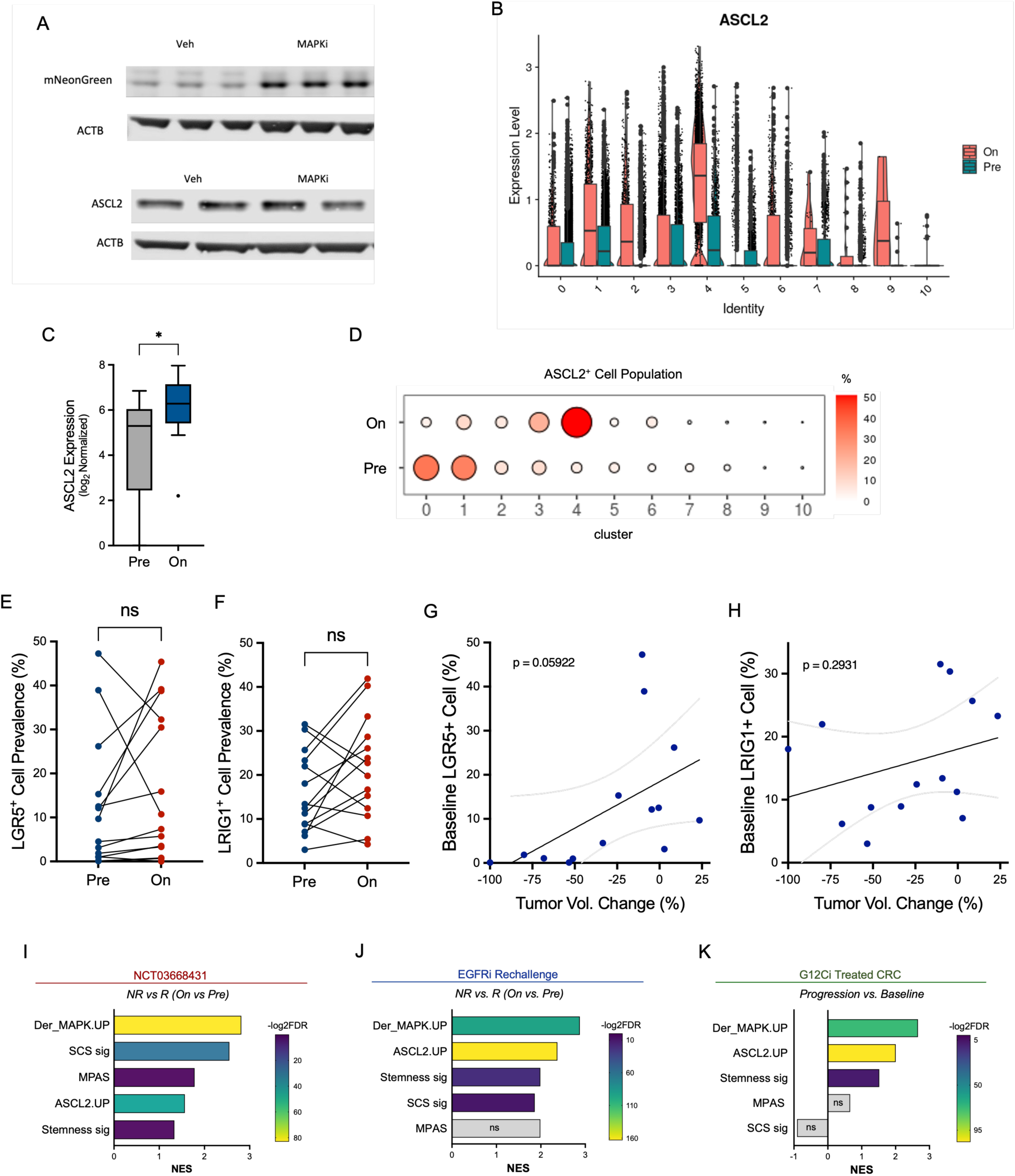
MAPKi increases an ASCL2 specific program. **A)** ASCL2 and GFP (mNeonGreen) western blots of CT26 ASCL2^GFP^ cells treated with vehicle or MAPKi therapy; ACTB as loading control, and samples ran as duplicate or triplicates. **B-H)** BRAFi patient cohort scRNAseq data results. **B)** Normalized ASCL2 expression in pre- (green) and on-treatment (pink) samples per cell cluster. **C)** Log normalized ASCL2 expression of cluster 4 cells in pre- and on-treatment samples, pseudobulked by patient (two-tailed Mann-Whitney). **D)** ASCL2^+^ cells denoted by ASCL2 gene expression, and prevalence of ASCL2^+^ population by cell cluster shown by timepoint (pre vs. on-treatment). **E-H)** Cells classified by LGR5 and LRIG1 gene expression, and LGR5^+^ (**E**) and LRIG1^+^ (**F**) cell prevalence in each patient (n = 14) sample compared between pre- (blue) and on- (red) treatment (two-sided paired t-test; LGR5: p = 0.32, LRIG1: p = 0.09). **G-H)** Pearson correlation using baseline (pre-treatment) LGR5^+^ (**G**) and LRIG1^+^ (**H**) cell prevalence compared to tumor volume change per patient; simple linear regression trendline and 95% CI. **I-K)** GSEA of MAPK and stem related signatures in NR versus R with MAPKi therapy, in the BRAFi patient cohort (**I**), the EGFRi rechallenge cohort (**J**), and (**K**) in patients (n=4) treated with G12Ci therapy at progression vs baseline.

**Supplementary Figure 7.**
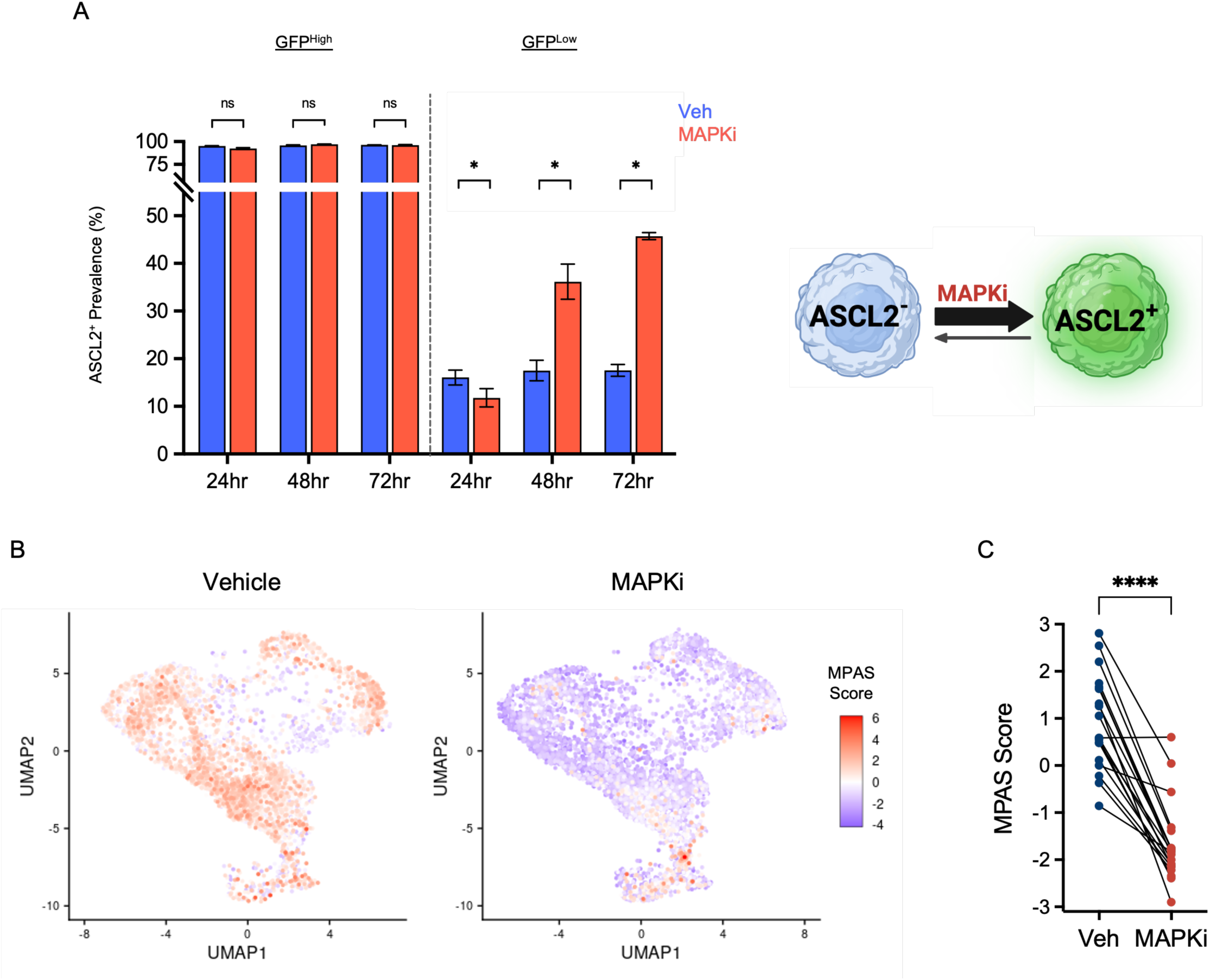
B1003b treatment suppressed MAPK activity. **A)** Flow-sorted B1003 ASCL2^GFP^ cells into GFP^High^ and GFP^Low^, and treated with vehicle (blue) or MAPKi therapy (red) for 72 hours, with samples (n = 3) re-analyzed by flow cytometry after treatment to determine ASCL2^+^ cell prevalence (two-way ANOVA with Bonferroni; mean ± SD); graphical illustration of MAPKi therapy shifting CRC cells into ASCL2^+^ state. **B)** UMAP of vehicle (left) and MAPKi (right) cells colored by their MAPK activity score (MPAS) expression. **C)** Quantification of MPAS expression in cells paired by matching barcodes across samples (two-sided paired t-test).

**Supplementary Figure 8.**
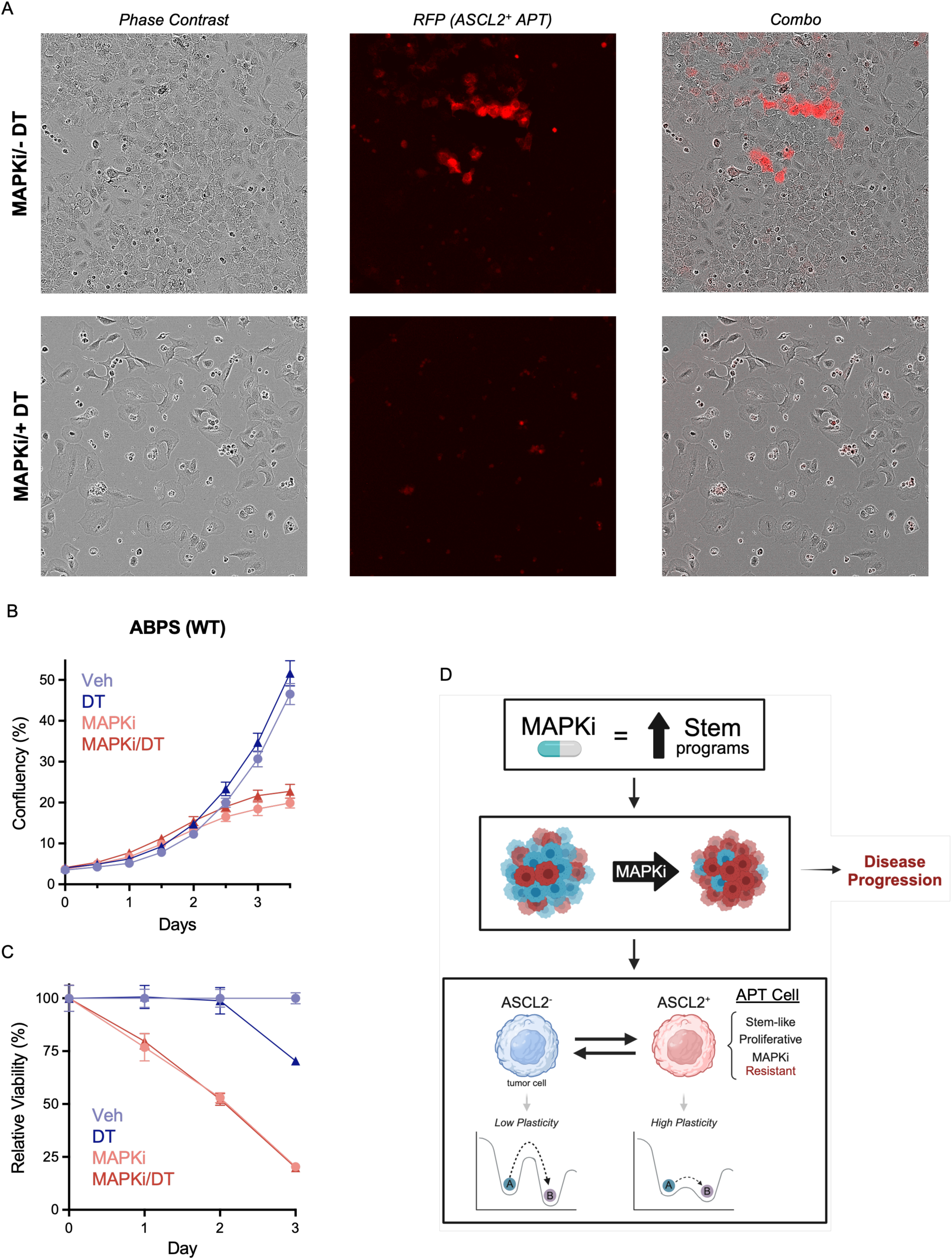
APT cells mitigate MAPKi response. **A)** Phase contrast, RFP, and overlapped images of CT26^DTR/RFP^ cells on day 10 of treatment with MAPKi/-DT or MAPKi/+DT. **B-C)** For all plots, the MAPKi treatment group is shown in red and vehicle group in blue. Data from samples treated with DT are triangles and vehicle (-DT) are circles. **B)** Growth assay of parental (WT) ABPS cells treated with vehicle, DT, MAPKi/-DT, or MAPKi/+DT; cell growth monitored by well confluency (n = 10; mean ± SEM). **C)** Relative cell viability, normalized to vehicle, of ABPS WT cells treated with vehicle, DT, MAPKi, or MAPKi/+DT for 72 hours (n = 6; mean ± SD). **D)** Summary of findings, outlining MAPKi therapy induces APT phenotype, enriching these cells in CRC tumors, and resulting in disease progression.

## Notes

### Competing Interest Statement

SK, ownership interest: Lutris, Iylon, Frontier Medicines, Xilis, Navire; consulting or advisory role: Genentech, EMD Serono, Merck, Holy Stone Healthcare, Novartis, Lilly, Boehringer Ingelheim, AstraZeneca/MedImmune, Bayer Health, Redx Pharma, Ipsen, HalioDx, Lutris, Jacobio, Pfizer, Repare Therapeutics, Inivata, GlaxoSmithKline, Jazz Pharmaceuticals, Iylon, Xilis, Abbvie, Amal Therapeutics, Gilead Sciences, Mirati Therapeutics, Flame Biosciences, Servier, Carina Biotech, Bicara Therapeutics, Endeavor BioMedicines, Numab, Johnson & Johnson/Janssen, Genomic Health, Frontier Medicines, Replimune, Taiho Pharmaceutical, Cardiff Oncology, Ono Pharmaceutical, Bristol-Myers Squibb-Medarex, Amgen, Tempus, Foundation Medicine, Harbinger Oncology, Inc., Takeda, CureTeq, Zentalis, Black Stone Therapeutics, NeoGenomics Laboratories, Accademia Nazionale Di Medicina, Tachyon Therapeutics; research funding: Sanofi, Biocartis, Guardant Health, Array BioPharma, Genentech/Roche, EMD Serono, MedImmune, Novartis, Amgen, Lilly, Daiichi Sankyo. All other authors have no competing interests to declare.

